# Jaccard dissimilarity in stochastic community models based on the species-independence assumption

**DOI:** 10.1101/2022.12.13.520233

**Authors:** Ryosuke Iritani, Vicente J. Ontiveros, David Alonso, José A. Capitán, William Godsoe, Shinichi Tatsumi

**Author notes:** Conception and design: RI (lead), VJO, DA, JAC, WG, and ST. Method development: RI. Acquisition of data: not applicable. Analysis: RI (lead), VJO, DA, JAC, WG, and ST. Interpretation: RI. Drafting of the first version: RI. Revision: RI (lead), VJO, DA, JAC, WG, and ST.

## Abstract

A fundamental problem in ecology is understanding the causes of change in species composition among sites (i.e. beta diversity). At present it is unclear how spatial heterogeneity in species occupancy across sites shapes these patterns. To address this question, we develop probabilistic models that consider two spatial or temporal sites, where presence probabilities vary both among species and between the sites. We derive analytical and approximate formulae for the expectation of pairwise beta-diversity. Using a novel graphical tool, Stochastic Incidence Plots (SIPs), which depict the presence probabilities in two sites across species labels, we develop a means to conceptualize the heterogeneity in presence probabilities: the steepness or unevenness of SIPs reflects species-level heterogeneity, while the degree of overlap between SIPs indicates site-level heterogeneity. Utilizing SIPs and a combinatorial approach in a two-species scenario, we demonstrate that beta-diversity is lower when SIPs are parallel compared to when they are anti-parallel. We also find that this prediction is testable with the well-known checkerboard pattern in incidence matrices. Finally, we applied the method to the species distribution models for five woodpecker species in Switzerland, showing that their spatial distributions will change significantly in the future. Overall, this work improves our understanding of how pairwise beta-diversity responds to occupancy heterogeneity.

## 1 Introduction

Beta-diversity represents the spatial or temporal variation in species compositions. Beta-diversity links diversity across scales (Whittaker 1972; Anderson *et al*. 2010; Chase *et al*. 2019; Poggiato *et al*. 2021), and varies with fundamental processes such as dispersal, environmental filtering and species interactions (Vellend 2010; Anderson *et al*. 2010; Socolar *et al*. 2016; Maynard *et al*. 2017; Legendre 2019; Thompson *et al*. 2020). The decline in beta-diversity, biotic homogenization, has raised significant concerns, as it may reduce ecosystem functioning across the globe (Olden & Poff 2003; Olden & Rooney 2006; Hautier *et al*. 2017; Mori *et al*. 2018; Olden *et al*. 2018; Albrecht *et al*. 2021; Wang *et al*. 2021; Rolls *et al*. 2023). Clarifying the patterns of changes in beta-diversity (‘beta-diversity patterns’) in response to variations in biotic and abiotic conditions should lead to improving our ability in biodiversity management, conservation, and urban planning in our modern society, from a spatially global perspective (Crowther *et al*. 2015).

Estimates for beta-diversity include pairwise indices derived from empirical presence-absence data (incidence data; Koleff *et al*. 2003). The incidence-based beta-diversity can take advantage of simplicity by focusing on binary data. Despite its simplicity, the patterns of variability in beta-diversity are not well conceptualized, with previous studies on incidence-based beta-diversity yielding mixed results. For example, some theoretical work shows that dispersal reduces beta-diversity (Loreau 2000; Thompson *et al*. 2020), whereas other shows that the impact of dispersal depends on metacommunity model-structure (Lu *et al*. 2019; Lu 2021). Meanwhile, experimental work suggests that dispersal may promote beta-diversity (Vannette & Fukami 2017). One of the challenging goals in the field is to make better sense of the complication in these beta-diversity patterns.

Organisms live in highly heterogeneous environments (Abson 2017), and such heterogeneity renders the problem of beta-diversity patterns even more challenging. That is, patterns of likely species occupancy (which we refer to as ‘chance of occurrence’ in this article) are highly heterogeneous. In the simplest scenario, the chance of occurrence is equal among all species across all sites (Figure 1A). This premise does not always hold. In general, species have different chances of occurrence across space, for instance due to differences in niches, exhibiting geographically interspersed distributions (Figure 1B; ‘checkerboard pattern’, Diamond 1975; Connor & Simberloff 1979). To provide another example, one site may have higher chance of occurrence for all species than other sites even if all species have an equal chance of occurrence (Figure 1C). In such cases, the incidence at one site is nested within that of another, thus contributing to large beta-diversity (Darlington Junior 1957; Patterson & Atmar 1986; Patterson 1987; Ulrich & Gotelli 2007). In general, chance of occurrence differs both among species and sites (Figure 1D), i.e., chance of occurrence is heterogeneous among species and across sites (‘occupancy heterogeneity’, MacKenzie *et al*. 2018, Chapter 4). The problem then is how we can better conceptualize beta-diversity patterns associated with occupancy heterogeneity.

**Figure 1:**
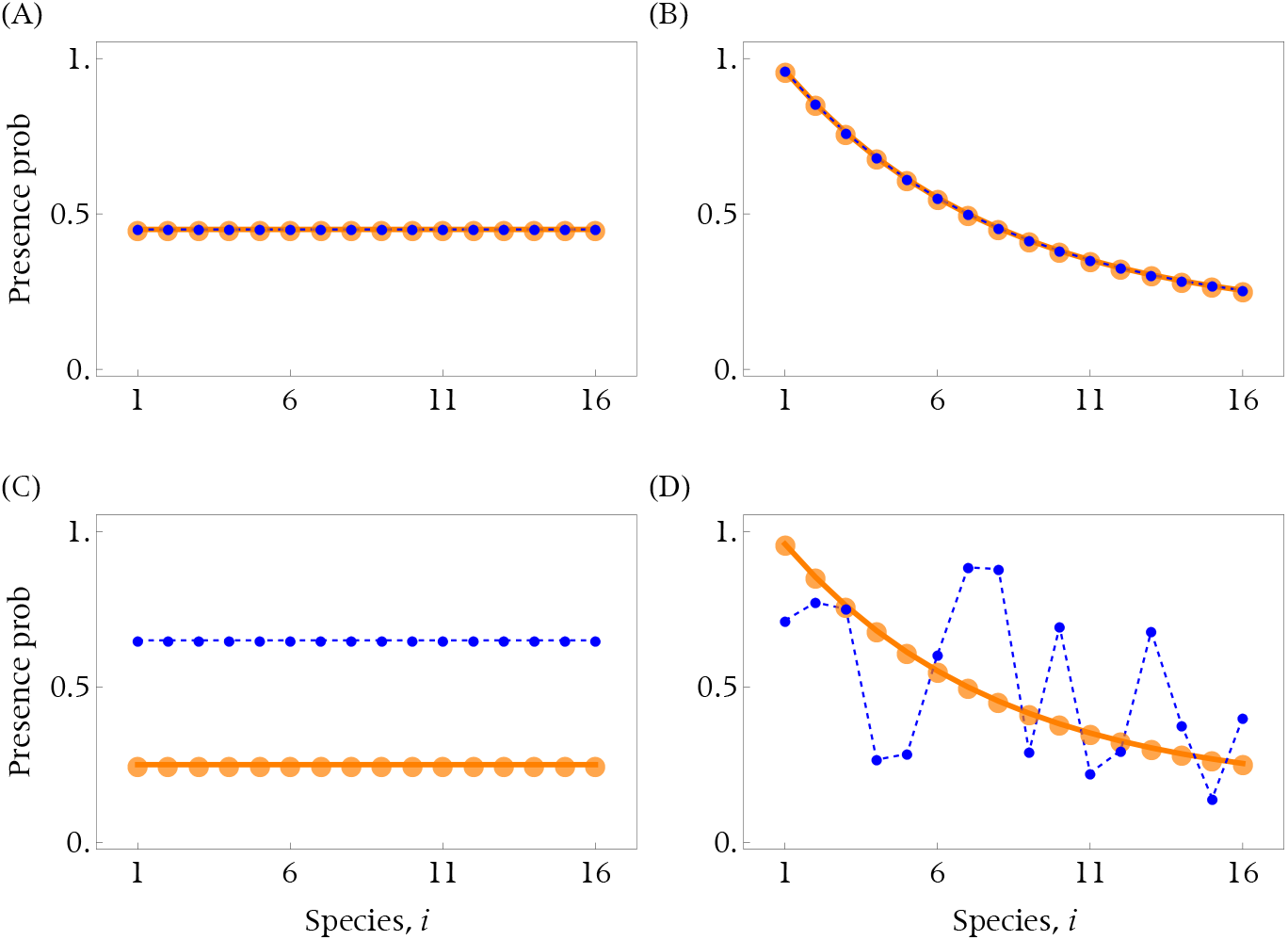
Stochastic Incidence Plots (SIPs). Presence probabilities in site 1 (blue) and site 2 (orange) are depicted along species labels. When two SIPs completely overlap, there is no heterogeneity between the two sites (A and B). When two SIPs are both flat, all species have the same chance of occurrence (A and C). In general, two SIPs are different and neither of them is flat (D), with numerous patterns of SIPs possible. SIPs therefore capture the intricate heterogeneity in the presence probabilities of species across space.

Theoretical models can help us predicting beta-diversity patterns. Recent work have developed a general model of metacommunity dynamics to study the effects of density-independent abiotic factors, density-dependent species interactions, and dispersal among patches on beta-diversity (Whittaker 1972; Thompson *et al*. 2020). However, Thompson *et al*. (2020) do not explicitly consider how beta-diversity co-varies with occupancy heterogeneity per se. To resolve this, we develop a general model where occupancy heterogeneity is modeled as the differences in the probabilities of species incidences. Applying the stochastic theory has several advantages. First, it allows us to study summary statistics (e.g., expectation and variance) of beta-diversity and how they vary with the variations in the presence probabilities among species and among sites, thus enabling to generate the null distribution of beta-diversity for hypothesis-making and testing (Gotelli & Graves 1996; Leibold *et al*. 2004; Gotelli & Ulrich 2011; Hui & McGeoch 2014; Chung *et al*. 2019; Thompson *et al*. 2020). Second, it allows us to consider various patterns of occupancy heterogeneity (Figure 1A–D), and therefore enables exploring the co-variability between beta-diversity and the heterogeneity. Third, the recent theoretical development of occupancy estimation is based on the stochasticity of species occurrences (MacKenzie *et al*. 2018).

Throughout the article, we assume that species incidences are independent of each other both within and between sites to nullify correlations among them, the so-called ‘species independence assumption’ (as in, e.g., Chung *et al*. 2019; Lu *et al*. 2019; Lu 2021; Kalyuzhny *et al*. 2021; Ontiveros *et al*. 2021). Biologically, the species-independence assumption implies that no biotic interaction occurs and environmental effects are captured by the probability that species are present. We derive the exact formulae, and approximation, of the expectation and variance of a pairwise beta-diversity index, Jaccard dissimilarity (Jaccard 1908, 1912; Veech 2012; Arita 2017; Keil *et al*. 2021). We then provide several theoretical predictions of the consequences of heterogeneity for the expectation of beta-diversity. In addition, we propose using a diagram (“Stochastic Incidence Plots”; SIPs) that depicts the presence probabilities along species labels for each site thereby easing a visual interpretation of heterogeneity (Figure 1). For example, non-flat (e.g., sloped) SIPs represent a scenario where presence probabilities differ among species; also, when two SIPs do not fully overlap (e.g., nested), two sites are differentiated in species presence probabilities. SIPs can therefore allow us to interpret the co-variability of beta-diversity and occupancy heterogeneity. We highlight the two-species case as the minimal model to fully analyze the relationship between beta-diversity and occupancy heterogeneity, by linking the SIPs and the probability distributions of incidence matrices. This analysis will be proven to be useful in allowing us to examine how a pair of species that have different chances of occurrence contributes to the increase or decrease in beta-diversity.

## 2 Method and Result

### Model

#### Incidence matrix

We consider a landscape consisting of two spatial locations (or temporal points) and compare the difference in species compositions. We write *S*_T_ for the maximum number of species present in the landscape, which we refer to as ‘species pool size.’

We write *X*_*r,s*_ for a binary state of incidence (*X*_*r,s*_ = 1 for presence and = 0 for absence respectively) of species *s* in site *r*, with *s* = 1, …, *S*_T_ and *r* = 1, 2. We generically write **X**, whose (*r, s*)-element is *X*_*r,s*_, for a 2-by-*S*_T_ incidence matrix.

#### Diversity

Here we describe diversity measures. The sum 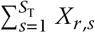 represents species richness in site *r*, and 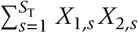 represents the number of commonly present species. Note that if species *s* is present in at least one of the sites, then *X*_1,*s*_ + *X*_2,*s*_ −*X*_1,*s*_ *X*_2,*s*_ = 1, otherwise 0. Therefore, gamma-diversity, defined as the number of species present in the landscape, is given by 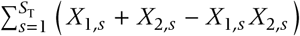. Similarly, if *s* is present in only one of the sites (referred to as being ‘unique’ in this literature), then *X*_1,*s*_ + *X*_2,*s*_ − 2*X*_1,*s*_ *X*_2,*s*_ = 1, otherwise 0. Then, the number of unique species is given by 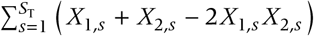.

Jaccard dissimilarity (JD) is then given by:

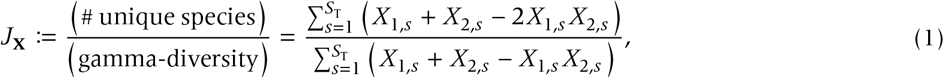

(Jaccard 1908, 1912; in Appendix A, we explain why JD is a suitable beta-diversity measure).

#### Stochastic analysis

Here we describe stochasticity in incidence matrices, assuming that species incidence is independent of each other (species independence assumption). We write *p*_*r,s*_ for the probability that species *s* is present in site *r*, i.e., probability that *X*_*r,s*_ = 1 (MacArthur & Wilson 1963; Real *et al*. 2016; Carmona & Pärtel 2020). For notational simplicity, we write *a*_*r,s*_ = 1 − *p*_*r,s*_ for the probability that species *s* is absent from site *r*. The species independence assumption implies that the probability of the occurrence of incidence matrix **X** is given by

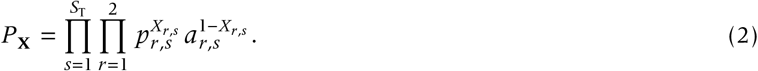

Next we define the following probabilities: species *s* is: (i) present in both sites with probability *b*_*s*_ := *p*_1,*s*_ *p*_2,*s*_ ; (ii) unique with probability *u*_*s*_ := *p*_1,*s*_ + *p*_2,*s*_ −2 *p*_1,*s*_ *p*_2,*s*_ ; or (iii) absent from both sites with probability *d*_*s*_ := (1 − *p*_1,*s*_) (1 − *p*_2,*s*_), with *b*_*s*_ + *u*_*s*_ + *d*_*s*_ = 1. Also, we write 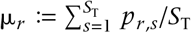 for the average presence probability in site *r*.

It is worth examining how these probabilities, *u*_*s*_, *b*_*s*_, and 1 − *d*_*s*_ = *b*_*s*_ + *u*_*s*_, change with presence probabilities *p*_1,*s*_, *p*_2,*s*_ (Figure S1). The probability of unique presence *u*_*s*_ is large when two sites have dissimilar presence probabilities (right bottom or top left regions). The probability of double-presence *b*_*s*_ is large when two sites have large presence probabilities. Finally, the probability of presence in at least one site 1 − *d*_*s*_ is large when two sites have large presence probabilities.

### Stochastic Incidence Plots (SIPs)

Given that beta-diversity varies with occupancy heterogeneity, how can we interpret such co-variability? We here introduce a graphical tool, Stochastic Incidence Plots (SIPs). SIPs consist of a pair of plots each depicting the presence probabilities in site 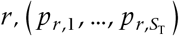 along species labels *s* = 1, …, *S*_T_, where we set *p*_1,1_ as the maximum among the all *p*_*r,s*_ ‘s.

SIPs provide a useful visual interpretation of occupancy heterogeneity (Figure 1). First, SIPs fully overlap each other when two sites have the same presence probabilities for each species (Figure 1A and B). Biologically, this scenario is relevant when: (i) presumably the environmental conditions in two sites are similar, or (ii) the actual environmental conditions are unknown and as such one uses a common value for both sites. Second, SIPs are both flat if and only if all species have the same presence probability in each site: 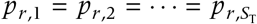 for each site, *r* = 1, 2 (Figure 1A and C). Biologically, this occurs when all species are equiprobably present (i.e., all species have the equal chance of occurrence). Finally, SIPs do not fully overlap nor are flat (Figure 1D), in which case there are diverse patterns of intersections of SIPs. SIPs can therefore allow us to qualitatively characterize the occupancy heterogeneity.

### Analytical result: Expectation of JD and its approximation

We denote the expectation of JD by E[*J* ] := ∑_**X**_ *P*_**X**_ *J*_**X**_ where the sum is taken over all possible incidence matrices. Using the moment-generating polynomials (Lange 2010), we get:

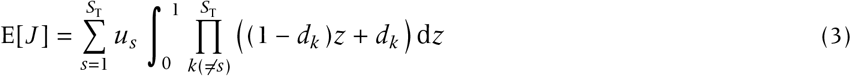

(Appendix B for derivation; note that we can calculate analytically the integral using the elementary polynomials and the beta-function). Eqn (3) reads as the sum of the probability that species *s* is present in only one of the sites (*u*_*s*_) multiplied by the integral of a polynomial product over an interval from 0 to 1 (with the polynomial referred to as probability-generating polynomial). In Appendix B, we prove that the integral for each *s* represents the reciprocal of the number of species (including species *s*) present in at least one of the sites. The conditional expectation, given that there is at least one species, is given by 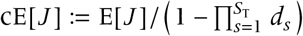.

We also derive an approximation of cE[*J* ] as:

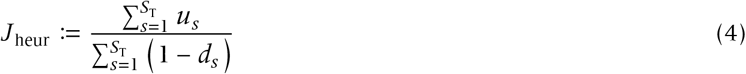

(Chung *et al*. 2019; Lu *et al*. 2019; Lu 2021; Kalyuzhny *et al*. 2021; Ontiveros *et al*. 2021; Appendix B), which we refer to as ‘heuristic approximation’ in this article. Eqn (4) reads as the expected number of unique species over the expected number of present species. We find that the heuristic approximation (Eqn (4)) gives a near-identical result with the conditional expectation (Figure 2), and converges to the exact value as the species pool size becomes large, *S*_T_ → ∞(Appendix B; one can also derive the exact and approximated variance of JD. For details, see Appendix B).

**Figure 2:**
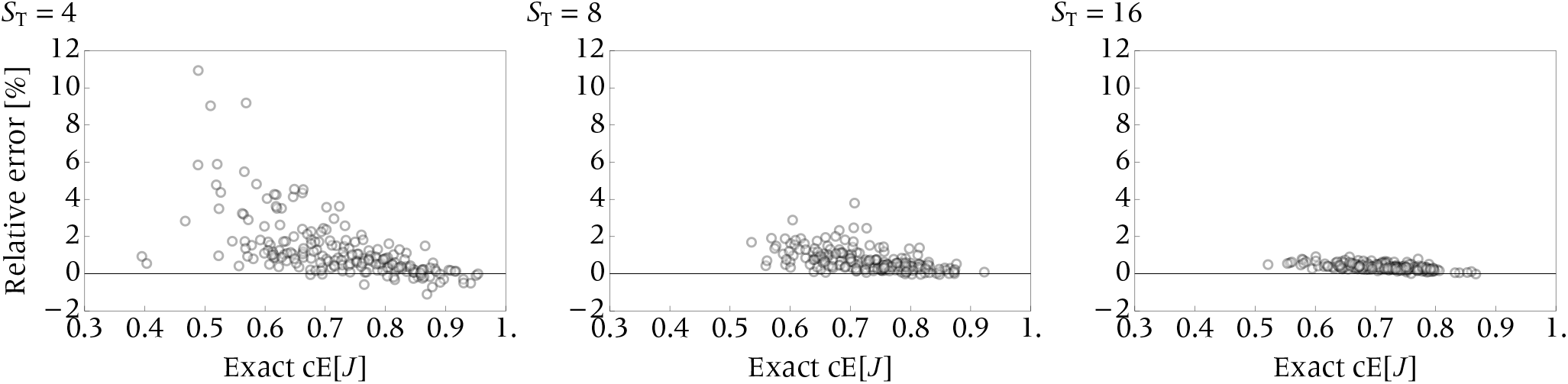
The relative error (defined as (*J* _heur_ − cE[*J* ])/cE[*J* ]) of the heuristic approximation, for different species pool sizes *S*_T_ = 4, 8, 16. We perform Monte-Carlo simulations by drawing *p*_*r,s*_ from the beta distribution (with parameters 0.95 and 1.15) and compute resulting relative errors. We run the stochastic simulations 200 times (thus 200 points in each panel). The heuristic approximation provides the near-identical approximation of the exact, conditional approximation.

### Benchmark result: species-equivalence (flat SIPs)

We here assume that all species have even chance of occurrence, so that SIPs are both flat (but not necessarily fully overlapping). In this case we find cE[*J* ] = *J* _heur_, that is, if *p*_*r,s*_ is common for all species for each *r* = 1, 2, then the heuristic approximation equates to the exact conditional expectation (Appendix C). Chung *et al*. (2019) and Ontiveros *et al*. (2021) have used the heuristic approximation for the special case where the presence probabilities are all equal (flat and overlapping; Figure 1A), and Lu *et al*. (2019) and Lu (2021) have generalized the result to the case where SIPs are flat but are not fully overlapping (Figure 1C). The present result thus generalizes the results of the previous literature.

### Key result 1: site-homogeneity (overlapping SIPs)

We next consider the scenario where SIPs fully overlap, while keeping the total average, μ_1_ + μ_2_, fixed. We show that as the chance of occurrence gets less even (i.e., more uneven), the expectation of beta-diversity gets smaller (Appendix C); specifically, the expectation is maximized when all species have the same chance of occurrence (Figure 3). We refer to this result as ‘transfer principle for beta’ (Chao & Ricotta 2019). Mathematically, the transfer-principle for beta states that the expectation of JD is a Schur-concave function (Tuomisto 2012; Marshall *et al*. 1979; Chao & Ricotta 2019). Graphically, flat, straight SIPs have the maximum beta-diversity than do sloped SIPs (Figure 3; see Appendix C for more details).

**Figure 3:**
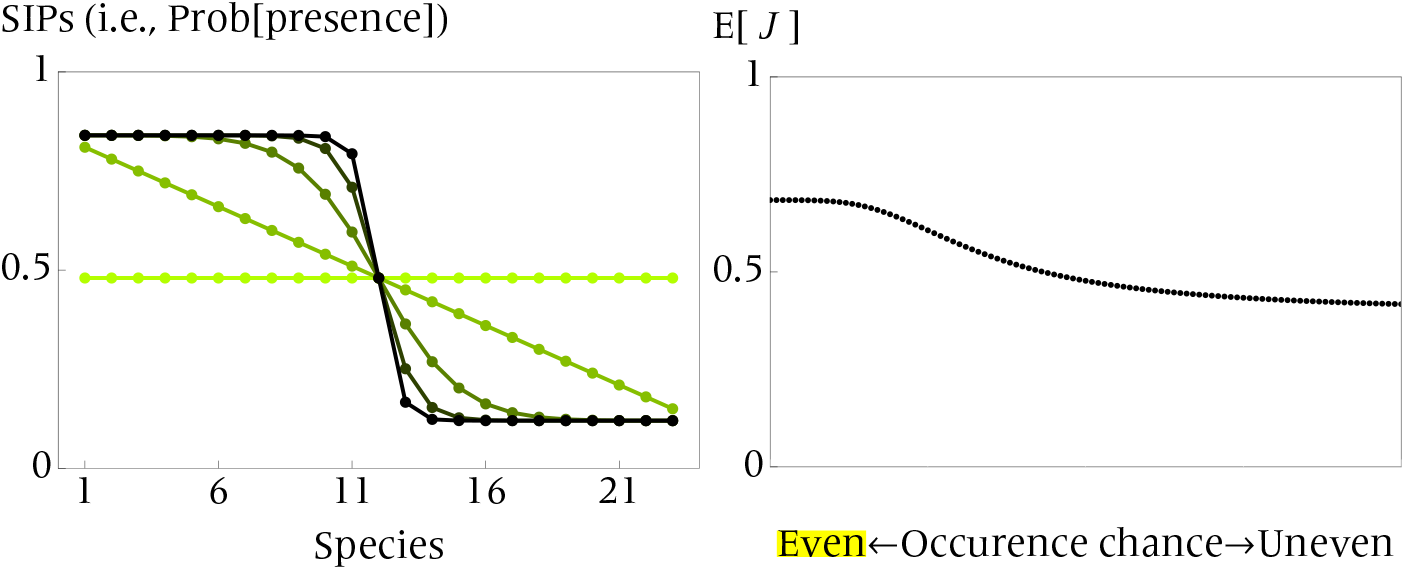
Key result 1, the transfer principle for beta. The presence probability is identical for both sites for each of the species out of *S*_T_ = 23 of species. As the chance of occurrence becomes uneven (i.e., SIPs become more sloped, from yellow to blue; left panel), the expectation of JD decreases (right panel). We use a Hill function that ranges from 0.12 to 0.84, with the average presence probability μ_1_ = μ_2_ = 0.48 kept fixed. An unevenness parameter is varied from 0 (yellow), 1, 4, 9 to 16 (= 4^2^ ; black). See Appendix C for more details.

### Key result 2: two species-case under heterogeneity

In the general case where species differ and sites are heterogeneous, it is hard to interpret co-variability of beta with the heterogeneity. We therefore focus on a case with two species *s* and *s*′, to facilitate intuitive understanding (note that here we use the alphabetical species labels *s* and *s*′ to avoid confusion with site labels 1 and 2). In this scenario, there are in total 2^2.2^ = 16 incidence matrices with different values of JD.

### Classification of incidence matrices

We first classify the 16 incidence matrices into seven types, based on the effect, on JD, of appending each matrix to a hypothetical incidence matrix. For example, suppose we have a hypothetical incidence matrix 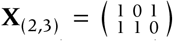 at hand; we append it a two-by-two incidence matrix 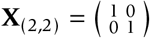 to get a new incidence matrix 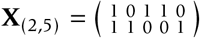. As 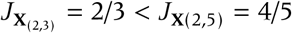, appending 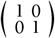 increases beta-diversity in this case.

We generalize this procedure. We examine the effects of appending each of 16 two-by-two incidence matrices to a given incidence matrix, and ask whether this appendix increases or decreases JD. As is summarized in Table 2, we find that: (i) three classes of matrices always increase JD; (ii) two of such three classes (named ‘nestedness’ and ‘checkerboard’ classes respectively) increase JD more than does the other; (iii) one class of matrices either increases or decreases JD, depending on the JD of the original, prepended matrix to be > 1/2 or < 1/2; (iv) there are two classes of matrices that always reduce JD, one of which (named ‘dominance’ class) reduces JD more than does the other; (v) JD is invariant with appending the all-zero matrix; and (vi) the incidence matrices other than nestedness, checkerboard, and dominance can be completely classified only by the average richness (marked ‘*’ in Table 2). We name such matrices in Table 2.

**Table 1:** Summary of notation used in the main text.

| Notation | Definition | Note |
| --- | --- | --- |
| $s$ | Species label | $s = 1, 2, \dots, S_T$ |
| $S_T$ | The total number of species in the mainland | “species pool size” |
| $r$ | Site label, with $r = 1$ or $2$ | “sites” may be spatial or temporal |
| $X_{r,s}$ | Incidence of species $s$ in site $r$ | 0 (absence) or 1 (presence) |
| $\mathbf{X}_{(2,S_T)}$ | Incidence table of size $S_T$-by-2 | Abbreviated to $\mathbf{X}$ |
| $:=$ | Defining a quantity | |
| $\mathbf{X}_{r,\circ}$ | Column vector of species incidences in site $r$ | |
| $\mathbf{X}_{\circ,s}$ | Row vector of species $s$'s incidence | |
| $p_{r,s}$ | Probability of $s$ present in $r$ | $a_{r,s} = 1 - p_{r,s}$ for probability of absence |
| $b_s$ | Probability of $s$ present in both sites 1 and 2 | b for “both” |
| $d_s$ | Probability of $s$ absent from both sites 1 and 2 | d for “double-absence” |
| $u_s$ | Probability of $s$ present in either site 1 or 2 | u for “unique” |
| $P_{\mathbf{X}}$ | Probability that a table $\mathbf{X}$ is observed | $= \prod_{s=1}^{S_T} \prod_{r=1}^2 p_{r,s}^{X_{r,s}} a_{r,s}^{1-X_{r,s}}$ |
| $J_{\mathbf{X}}$ | Jaccard dissimilarity for an incidence table $\mathbf{X}$ | |
| $\gamma_{\mathbf{X}}$ | The number of species present in the landscape for table $\mathbf{X}$ | “Gamma-diversity” |
| $E[J]$ | Expectation of Jaccard dissimilarity | $cE[J]$ for conditional expectation |
| $J_{\text{heur}}$ | Approximation of $cE[J]$ | “Heuristic approximation” |
| $\mu_r$ | Average presence probability in site $r$ | $= \sum_{s=1}^{S_T} p_{r,s} / S_T$ |

**Table 2:** Classification of incidence matrices. Seven classes characterize two-by-two matrices. The incidence matrices that have the average number of species (alpha-diversity, α) per site being 1 are further subclassified according to nestedness, checkerboard, and dominance patterns. The name ‘dominance’ accounts for the fact that in both sites, one species has higher chance of occurrence than does the other. Otherwise if α = 0, 0.5, 1.5, or 2, then each of them consists a single class. Symbol ‘++’ indicates it has larger effect than does ‘+’, and ‘−−’ indicates it has a negatively larger effect than does ‘−’. The plus-minus symbol (‘±’) indicates a mixed effect (either increases or decreases JD).

| Incidence class, $C$ | Characteristics | Effect on beta |
| --- | --- | --- |
| $\begin{pmatrix} 1 & 1 \\ 0 & 0 \end{pmatrix}, \begin{pmatrix} 0 & 0 \\ 1 & 1 \end{pmatrix}$ | nestedness | ++ |
| $\begin{pmatrix} 1 & 0 \\ 0 & 1 \end{pmatrix}, \begin{pmatrix} 0 & 1 \\ 1 & 0 \end{pmatrix}$ | checkerboard | ++ |
| $\begin{pmatrix} 1 & 0 \\ 0 & 0 \end{pmatrix}, \begin{pmatrix} 0 & 1 \\ 0 & 0 \end{pmatrix}, \begin{pmatrix} 0 & 0 \\ 1 & 0 \end{pmatrix}, \begin{pmatrix} 0 & 0 \\ 0 & 1 \end{pmatrix}$ | $\alpha = 0.5^*$ (single-one) | + |
| $\begin{pmatrix} 1 & 1 \\ 1 & 0 \end{pmatrix}, \begin{pmatrix} 1 & 1 \\ 0 & 1 \end{pmatrix}, \begin{pmatrix} 1 & 0 \\ 1 & 1 \end{pmatrix}, \begin{pmatrix} 0 & 1 \\ 1 & 1 \end{pmatrix}$ | $\alpha = 1.5^*$ (single-zero) | ± |
| $\begin{pmatrix} 1 & 0 \\ 1 & 0 \end{pmatrix}, \begin{pmatrix} 0 & 1 \\ 0 & 1 \end{pmatrix}$ | dominance | - |
| $\begin{pmatrix} 1 & 1 \\ 1 & 1 \end{pmatrix}$ | $\alpha = 2^*$ (all-one) | -- |
| $\begin{pmatrix} 0 & 0 \\ 0 & 0 \end{pmatrix}$ | $\alpha = 0^*$ (all-zero) | 0 |

### SIPs by permutation: fundamental SIPs

To consider numerous shapes of SIPs, we use a permutation approach. We pick four (generally distinct) arbitrary values between 0 and 1, and assign their maximum to *p*_1,*s*_, the presence probability of species *s* in site 1 (without loss of generality). We then assign the remaining three to the other presence probabilities *p*_1,*s*_′, *p*_2,*s*_, *p*_2,*s*_′, obtaining 3! = 6 pairs of SIPs (Figure 4). We refer to these six pairs of SIPs as ‘fundamental SIPs.’

**Figure 4:**
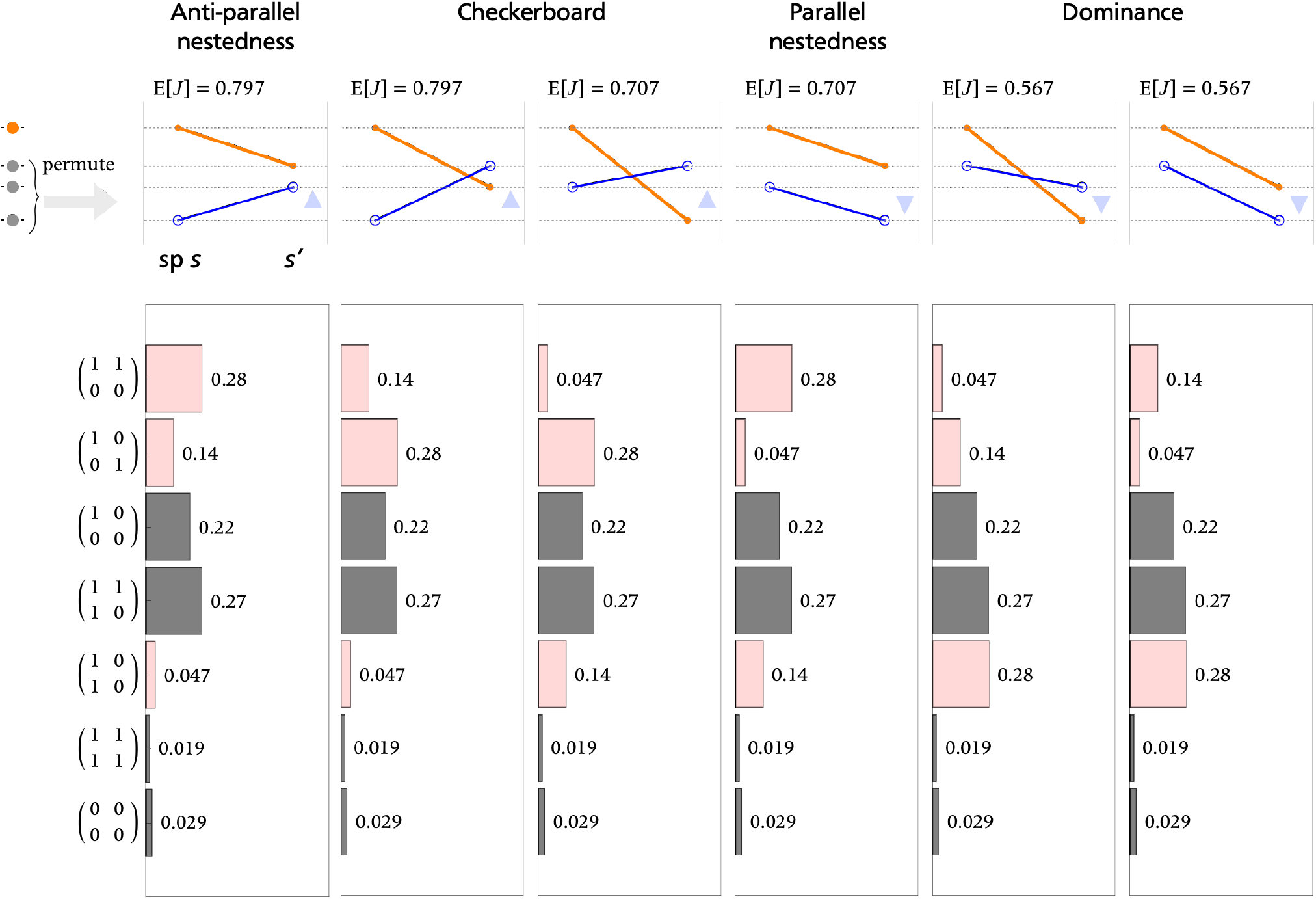
The permutation approach to produce the six fundamental SIPs (top six panels), with four values (0.88, 0.56, 0.38, and 0.10) assigned to the presence probabilities. The histograms of the seven classes of incidence matrices, in which gray bars are common across all the fundamental SIPs, whereas the light red bars are not. Anti-parallel nestedness SIPs most likely produce the nestedness class (top bar). Checkerboard SIPs most likely produce the checkerboard class (second top bar). The parallel nestedness SIPs also most likely produce the nestedness class, but also likely produce the dominance class (fifth bar) that reduced JD. The dominance SIPs most likely produce the dominance class (third bottom bar). Blue triangle indicates a slope of SIP in site 2 (upward or downward).

The advantage of using this permutation approach is twofold. First, by construction, the fundamental SIPs exhaustively cover possible shapes of SIPs, including crossed, parallel, and anti-parallel ones. Second, we can fix the total average of presence probability (μ_1_ + μ_2_)/2 kept constant, so that we can isolate the effect of heterogeneity on the expectation of JD, regardless of the average presence probabilities per se.

We then calculate the expectation of JD for each of the fundamental SIPs. We obtain three (= 6/2) types of expected values of JD, as JD is invariant with permutation of site labels for each species (i.e., swapping *p*_*r*,1_ with *p*_*r*,2_ does not change the expectation of JD).

We find that three pairs of fundamental SIPs are crossed (each with different expectations of JD) and the others are uncrossed (Figure 4; Appendix C). Among the uncrossed pairs of fundamental SIPs, two of them are ‘parallel’ (both SIPs cline downward) and the remaining are ‘anti-parallel’ (one of the SIPs upward while the other downward). In addition, we find that the anti-parallel pair tends to have larger beta-diversity than does the parallel pair.

We categorize the six types of fundamental SIPs into four types (Figure 4): (1) Anti-parallel nestedness, where SIPs are anti-parallel but uncrossed; (2) Checkerboard, where both SIPs are parallel and crossed; (3) Parallel nestedness, where SIPs are parallel and uncrossed, and (4) Dominance, where one species’ minimum presence probability exceeds the other’s maximum.

### Relation between the seven classes and six SIPs

We now ask which of the fundamental SIPs contributes to larger or smaller expected values of JD, by examining how likely each of the six fundamental SIPs produces which class of matrices (Figure 4). Towards this end, we generate a histogram (probability distribution) of the seven classes generated by each of the fundamental SIPs, using:

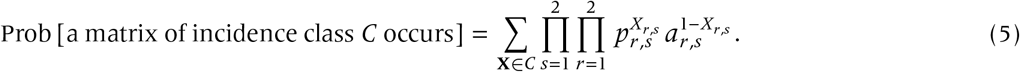

We find that, with the same probability, the fundamental SIPs produce each of the single-one, single-zero, all-one, and all-zero matrices (Figure 4 gray bars,Table 2; Appendix C). As such, with different probabilities, the other classes (nestedness, checkerboard, and dominance classes) are generated by the fundamental SIPs (Figure 4 red bars).

To clarify the difference among the nestedness, checkerboard, and dominance incidence matrices: we derive the following findings from the histogram (but also based on mathematical arguments; see Appendix C): the anti-parallel nestedness SIPs most likely produce the nestedness incidence matrices; the checkerboard SIPs most likely produce the checkerboard incidence matrices; the parallel nestedness SIPs most likely (as likely as do the anti-parallel nestedness SIPs) produce the checkerboard incidence matrices; and the dominance SIPs most likely produce the dominance incidence matrices (Appendix C). We can generalize this result for any four distinct values of presence probabilities we use to produce the fundamental SIPs (Appendix C). Hence, we can characterize the relationship between the fundamental SIPs and the incidence classes.

To gain a better insight of beta-diversity patterns, we compare the difference in the expected values of the fundamental SIPs. Specifically, we examine the expectation of JD for the parallel versus anti-parallel pairs of fundamental SIPs. We find that anti-parallel pairs have larger JD than does the parallel pairs. This prediction can be understood in light of the association between the SIPs and incidence matrices: the parallel pair corresponds to dominance incidence matrices (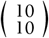 and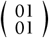), in which one species dominates over the other in the occupancy in both sites, thus reducing JD. For the same reason, the anti-parallel pair more likely generates checkerboard patterns than does the parallel pair, resulting in larger JD. Taken together, these findings provide clear interpretations of the relation among SIPs, incidence matrices, heterogeneity, and beta-diversity, that is, the beta-diversity patterns associated with occupancy heterogeneity.

### Application to species distribution model

Applications of the present method encompass species distribution models (SDMs; Elith & Leathwick 2009; Guisan *et al*. 2017; Zurell *et al*. 2020). Generally, SDMs seek to estimate the presence probability of each species at a spatial site by using environmental information (e.g., temperature). As SDMs assume that the presence probability of each species is independent of each other, SDMs are well suited to the assumption of the present method. We apply the present method to SDMs of five woodpecker species (*Picus viridis, P. canus, Dendrocopos major, D. minor*, and *Dryocopus martius*; Appendix D), using the temporal analogue of beta-diversity (temporal beta-diversity) to estimate the difference in the present and future spatial distributions of species (Legendre 2019; Magurran *et al*. 2019). We find that the expectations of temporal JD are high across the landscape (Figure 5), suggesting that the distribution of the woodpeckers will change significantly in the future. We also find that this result is primarily explained by species dynamics in lowland sites where some species thrived and others failed: *P. canus*, that will decrease its occupancy rate near the rivers and will increase in surrounding areas (Figure 5), and *D. minor*, whose occupancy is expected to increase in lowlands and valleys (Figure 5). Dissimilarity in hillsides is expected to be moderate due to a general increment in richness (Figure 5). These results are consistent with a general trend of Switzerland forest birds moving to higher grounds as a response to environmental change (Maggini *et al*. 2014). See Appendix D for the full details.

**Figure 5:**
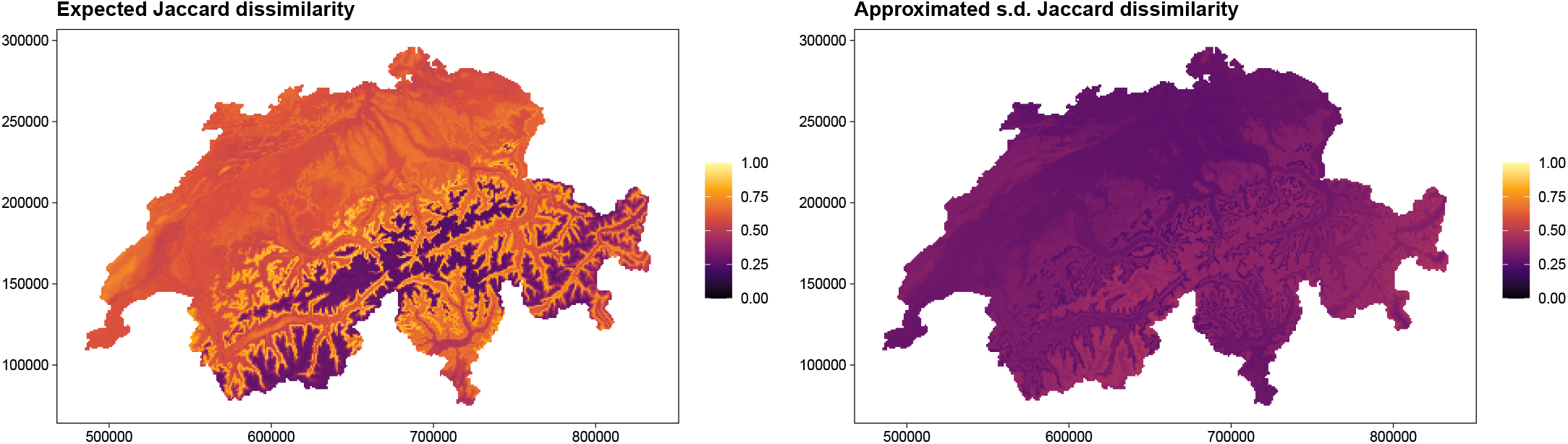
The change in the distributions of woodpeckers. We have quantified the expected, compositional dissimilarity of five woodpecker species at two time points, current and future, over the region of Switzerland. We have used occupancy estimations for current and future climatic conditions over Switzerland. (A) Expectation. Compositional changes are expected to be high in the upper limit of the current distribution and lowlands. (B) Standard deviation (approximated, using Eqn (B71)). The standard deviation is overall small.

## 3 Discussion

Understanding the patterns of the variability in beta-diversity estimates in response to heterogeneity is of crucial importance in ecology. We studied the effect of heterogeneity in species’ presence probabilities (occupancy heterogeneity) on beta-diversity (Figure 1). We derived the analytical and approximated formula of the expectation of Jaccard dissimilarity index (JD) assuming that presence probabilities of species across sites are generally different (Figure 2). We first found that, when all species have the same chance of occurrence in each site, the formula of the expectation of JD used in previous studies provide the analytical expression of the conditional expectation of JD provided that at least one species is present (Chung *et al*. 2019; Lu *et al*. 2019; Lu 2021; Kalyuzhny *et al*. 2021; Ontiveros *et al*. 2021). Using the formula and numerical approaches, we found that when each species has the equal chance of occurrence in two sites, the evenness in the chance of occurrence yields larger expectation of JD than does the uneven case (the transfer principle for beta; Figure 3). Also, using the graphical approach, which depicts the presence probabilities along species for each site (stochastic incidence plots, SIPs), we classified the patterns of SIPs associated with the incidence matrices (e.g., checkerboard and nestedness; Table 2). We identified two types of SIPs that produce the large expectation of JD: (a) the anti-paralleled nestedness SIPs, in which two SIPs are anti-parallel and one site is nested by the other, and (b) the checkerboard SIPs, in which two SIPs have opposite slopes (upward and downward) and crossed. In contrast, we also found that if there is dominance relationship between species, in which a portion of species have larger chances of occurrence than do others, then the expectation of JD tends to be low (Figure 4).

We found that when each species has the same chance of occurrence for two sites, or that is when two sites are homogeneous, if the chances of occurrence are more even among species, then the expectation of JD is larger (transfer principle for beta). This prediction suggests that if certain species are dominant in the chance of occurrence over others, then the beta-diversity tends to be small. Consistently, the two-species scenario where the anti-parallel nestedness SIPs have larger beta-diversity than does the parallel nestedness SIPs; in the anti-parallel SIPs, the species’ chances of occurrence are balanced between sites, whereas the parallel SIPs have certain species have dominantly higher chance of occurrence over the other. This consistency suggests that chance of occurrence is balanced among species when the beta-diversity is large. Also from a perspective of beta-diversity estimation, the transfer principle for beta suggests that using a common, averaged presence probability for all species (flat SIPs) may result in systematic overestimation of beta-diversity. From a conservation biological perspective, the differences in species’ chance of occurrence alone would yield, counter-intuitively, lower spatial beta-diversity when spatial environment is homogeneous, suggesting that preserving spatial heterogeneity is needed to retain large beta-diversity. Taken together, the present result has various implications for beta-diversity patterns associated with occupancy heterogeneity.

We carried out a detailed analysis of the relationship between occupancy heterogeneity and the expectation of JD for the simplest, two-species case. We identified the patterns of SIPs associated with the probability of certain incidence matrices to occur. We categorized the SIPs, which we generated by a permutation of four distinct probability values, into four types: anti-parallel nestedness, checkerboard, parallel nestedness, and dominance. These categories have directly to do with empirically suggested patterns of incidence matrices. The anti-parallel nestedness SIPs most likely produce nested patterns of incidence matrices, whereas the checkerboard SIPs most likely produce the checkerboard patterns of incidence matrices. The parallel nestedness SIPs also most likely produce the nested patterns of incidence matrices, but have a lower expectation of JD because these SIPs also likely to produce dominance patterns of incidence matrices. Dominance SIPs likely yield a low expectation of JD because when one species is more likely to be present in both sites than does the other, it is likely that the incidence matrices have mixed or negative effects on JD. From this categorization, we can gain better insights on the estimates of beta-diversity, especially in relation to occupancy heterogeneity.

We showed that the heuristic approximation of JD agrees well with the conditional expectation of JD and its accuracy improves as the species pool size increases. The heuristic approximation is computationally much cheaper than calculating the exact in a brute force approach by avoiding integrating the polynomial products. We therefore suggest that when species pool size is large, using the heuristic approximation is suitable. If one wishes to evaluate the exact expectation, applying Gauß’ fast Fourier Transforms can be a good solution to improve the computational speed (Cooley & Tukey 1965; Heideman *et al*. 1984). We remark that the heuristic approximation uses the average values of uniqueness and double-presence probabilities, not the average values of the presence probabilities. Using the averaged presence probabilities corresponds to asking whether flat SIPs yield a good approximation of generally non-flat SIPs, and we found that the relative error can be large, although it becomes smaller as the species pool size increases (Appendix B). By considering all these facts, researchers may decide which of the statistics of JD to use.

We used a parsimonious approach, assuming the minimum species pool size *S*_T_ = 2, to examine the relationship between SIPs and incidence matrices. This approach allows for a generalization to larger species pool sizes. For example, even if the species pool size is large like *S*_T_ = 100, we can extract a subset pair of species *s, s*′, and examine whether the pair of species have nestedness, checkerboard, or dominance relationship. This approach has long been used in ecology, with a history dating back to the early studies of community assemblages and island biogeography (MacArthur & Wilson 1963; MacArthur & Wilson 1967; Diamond 1975; Connor & Simberloff 1979), nowadays called as ‘null model analysis’ (Gotelli & Graves 1996; Gotelli 2000; Peres-Neto *et al*. 2001; Leibold *et al*. 2004; Gotelli & Ulrich 2011; Hui & McGeoch 2014). Building up the theoretical framework from the simplest to more complex (and realistic) scenarios could be a next step to advance the field of quantitative ecology.

Also using the two-species case, we showed that the anti-parallel nestedness SIPs have larger beta-diversity than do the parallel SIPs. In the anti-parallel nestedness SIPs (one SIP upward and the other downward) that have a larger beta-diversity than do the parallel SIPs (downward SIPs), one species has a larger chance of occurrence in a site but a lower chance of occurrence in the other site, than does the other species, which means that these species tend to shape the checkerboard incidence patterns and thus have a large beta-diversity. In contrast, in the parallel SIPs, one species dominates over the other in the chance of occurrence, likely shaping the dominance incidence patterns (one species is found commonly but the other is totally absent). In addition, as the steepness of parallel nestedness SIPs gets larger, the likeliness of dominance matrices becomes larger and therefore the expectation of JD gets smaller. On the other hand, as the steepness of anti-parallel nestedness SIPs gets large, the likeliness of checkerboard matrices becomes larger and therefore the expectation of JD gets larger. As these incidence matrices have been the most prominent patterns in ecology assemblage theory (Diamond 1975; Connor & Simberloff 1979), the present framework suggests that it is promising to explicitly incorporate occupancy heterogeneity into existing theory to better understand beta-diversity patterns. Comparing SIPs with specific incidence matrices can offer insights into spatial incidence patterns, and using this framework may enhance our understanding of heterogeneity’s impact on beta-diversity.

Applying the method to species distribution models (SDMs) shows that woodpecker distributions in Switzerland will change significantly in the future. Likely mechanisms of the species differences and temporal heterogeneity in this system include colonization abilities, habitat selection, and species-specific tolerance to environmental challenges. In our worked example, we identify areas of interest for monitoring or management actions where assemblage composition will change under future climate conditions. Our method is therefore valuable for conserving and managing species distribution shifts caused presumably by ongoing global change.

Our study has implications for conservation. Generally, beta-diversity is a key factor for ecosystem functioning from local to global scales (Socolar *et al*. 2016; Mori *et al*. 2018). Local ecosystem functioning may be driven by species’ functional dissimilarity like niches (Godoy *et al*. 2020). For example, Loiseau *et al*. (2016) pointed out that conservation management designed to protect taxonomic diversity cannot be fully reconciled with functional diversity management. This result has to do with our prediction that, when two sites are fully homogeneous, the uneven chance of occurrence reduces beta-diversity. Therefore, considering that species’ difference in chance of occurrence is correlated with functional diversity (Palacio *et al*. 2022), where environmental similarity results in the similarity of presence probabilities of functionally similar species, the present prediction suggests that a conservation management aiming to maintain high beta-diversity be traded-off against the local, functional diversity. This trade-off becomes more complex when the two sites are heterogeneous in species’ chances of occurrence. One promising approach to advance the field is thus to identify the condition for the chance of occurrence of species to be balanced (i.e., even). This then produces new interesting questions. Moving forward, open questions include: how does incidence-based beta-diversity respond to changes in functional diversity in colonization ability and extinction tolerance (the capabilities of persistence against disturbances)? How does functional diversity, in turn, respond against the reduction in compositional dissimilarity (biotic homogeneization)?

To conclude, we investigated how beta-diversity varies with the heterogeneity in the chances of occurrence of species in two sites. We analyzed the expectation and variance of Jaccard dissimilarity index. Using a graphical representation, SIPs, we provided several key predictions for the effect of occupancy heterogeneity on beta-diversity. This work will help researchers better understand the probabilistic, or stochastic, nature of Jaccard dissimilarity (Real & Vargas 1996). Future studies may explore the effects of species associations on the probabilistic properties of Jaccard dissimilarity, and also carry out occupancy dynamics analyses, beyond pairwise dissimilarity analyses (MacKenzie *et al*. 2018). A promising approach includes a process-based approach (Thompson *et al*. 2020; Pilowsky *et al*. 2022), by which we can incorporate further complications that influence beta-diversity. Our method can incorporate additional realities to track and manage the changes in species distributions under global changes.

## Acknowledgement

The authors thank Ryosuke Nakadai, Naoto Shinohara, and Akira Terui, for helpful comments. We thank JSPS-KAKENHI (grant numbers 19K22457, 19K23768, 20K15882, 24H02291, and 24H01528 to RI, and 21K14880 to ST) for funding. We also thank Margarita Salas grant funded by the Spanish Ministry of Universities and the “European Union - Next GenerationEU” to VJO, CRISIS (PGC2018-096577-B-I00) to DA and JAC, UNIQUE (PID2021-127202NB-C21) to DA, and PRIORITY (PID2021-127202NB-C22) to JAC, all funded by MCIN/AEI/10.13039/501100011033 and “ERDF A way of making Europe”. WG was supported by the Te Pūnaha Matatini centre of research excellence. RI was inspired by some of the questions and answers on Cross Validated (Stack Exchange), in calculating the expectation of reciprocals, and also highlight a pioneering paper Cressie *et al*. (1981) on the calculation of expectations of reciprocals.

## Notation

We use the following notation throughout the appendix.

° Ω := {0, 1}
° *S*_T_ : The species pool size, *S* := # {*s* | max_*r*=1,2_ *p*_*r,s*_ > 0}
° *X*_*r,s*_ ∈ Ω: Incidence matrix
° 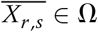: Logical negation, i.e., 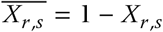
° *p*_*r,s*_ : Probability that *X*_*r,s*_ = 1
° *a*_*r,s*_ : Probability that *X*_*r,s*_ = 0
° **X** ∈ Ω^2^ ⊗ Ω^*S*^T : Incidence table of size with 2 rows and *S*_T_ columns.
° 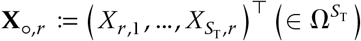
° *P*_**X**_: Probability that the incidence table **X** realizes
° 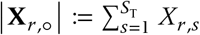 The total number of species present in a site *r*
° **X = (X**_**1**_**°, X**_**2**,_ **°**) ^⊤^ as we consider only two sites.

## Appendix A Jaccard dissimilarity

### Definition of Jaccard dissimilarity

We write *J*_**X**_ for Jaccard dissimilarity (JD) index for a table **X**, defined by:

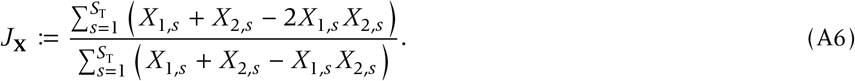

For **X** = **O** (zero-matrix), we define *J*_**O**_ := 0, which follows from two facts: (i) two all-zero vectors are (or axiomatically should be) completely similar, and (ii) the nullification of the denominator (which is always larger or equal to the numerator) should imply the nullification of the numerator (which is smaller or at most equal) as well. To avoid confusion, we suppose that numerator being zero implies JD be zero (otherwise resulting in erroneous calculations). It makes sense to exclude the zero-matrix, because zero-matrix indicates that there is no species in the landscape. Therefore we will focus on the conditional expectation.

### Linking JD to Whittaker’s (1972) beta-diversity

Whittaker (1972) defined beta-diversity as the ratio between region-wide diversity (gamma) to average local richness (alpha). Because the average local richness is 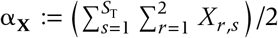, we have:

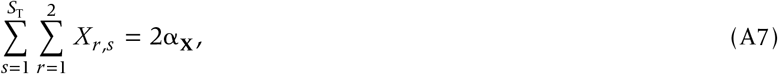

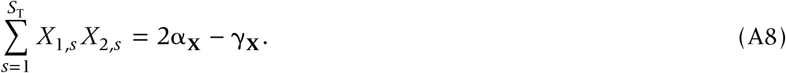

Using the beta-diversity sensu Whittaker (1972), which is given by:

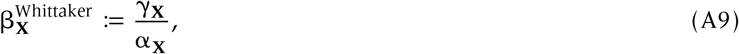

JD can be rewritten as:

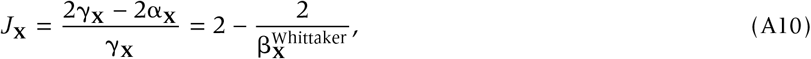

giving:

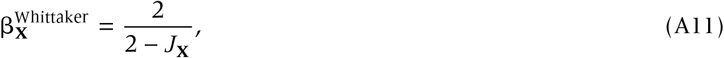

which is a monotonically increasing function of *J*_**X**_. Geometrically, 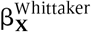, is a slope between two points (2, 2) an (*J*_X_, 0) which becomes steeper as *J*_**X**_ increases. Because Whittaker’s (1972) beta-diversity is a monotonic transform of JD, we can use JD as a measure of beta-diversity.

## Appendix B Stochastic analysis

**Figure S1:**
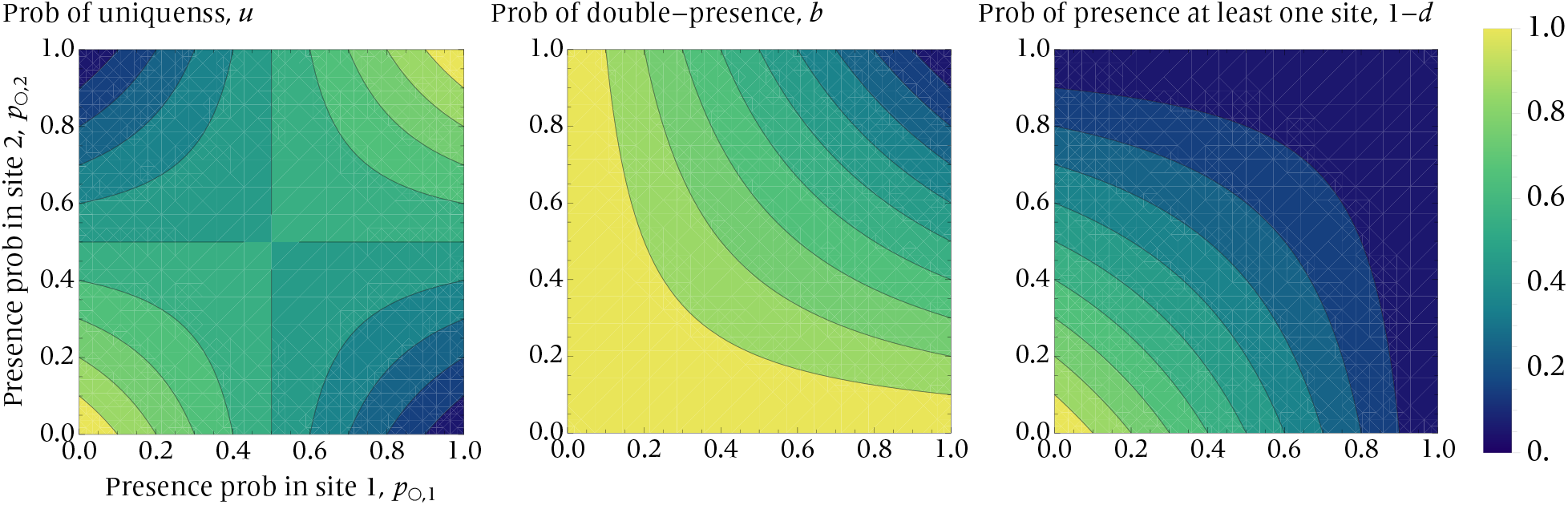
The dependence of he probabilities of uniqueness, double-presence, and presence in at least one site, on the pair of presence probabilities.

### Expectation

#### Step 1: express JD as an integral

We use the following identity:

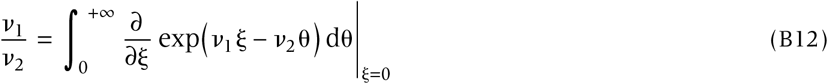

for any nonnegative values *v*_1_ and *v*_2_, which yields:

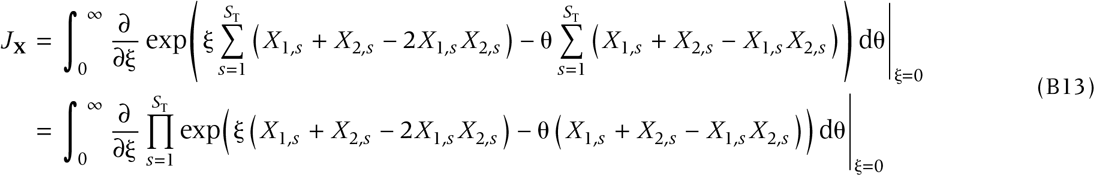

where we assign that we do not interchange the integral with the derivative unless otherwise stated, in order to remind that the integral should be defined as zero whenever the numerator is zero. We compute the expectation of *J*_**X**_ (which is a stochastic variable) over the distribution *P*_**X**_.

#### Step 2: Independence yields product

Assuming the species independence, the probability that a given incidence table **X** occurs is given by:

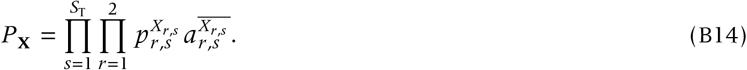

Then we get the (unconditional) expectation of JD as:

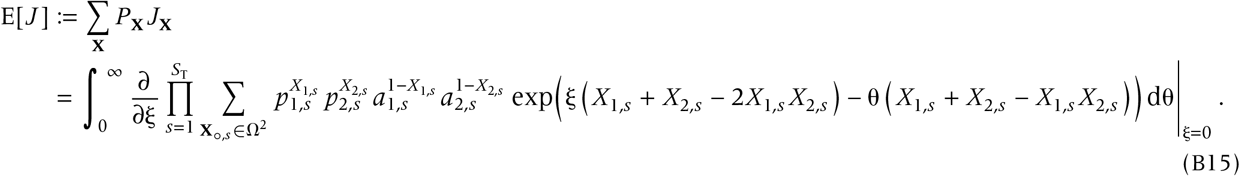

#### Step 3: Boolean thinking

Let us evaluate the Boolean variable:

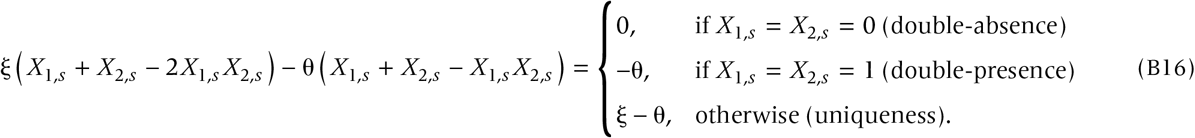

Using this allows us to expand the summation 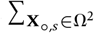 to get:

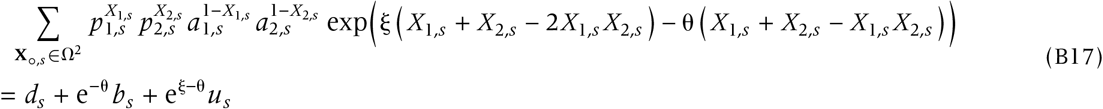

for all *s* ∈ {1, …, *S*_T_}. Therefore, substituting this into Eqn (B15) results in:

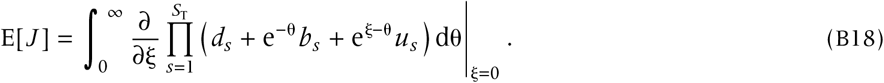

#### Step 4: apply Leibniz rule

By using Leibniz rule of the derivative of a product, we can get:

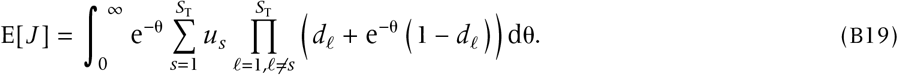

By transforming the variable *z* = 1 − e^−θ^ with dθ = (1 − *z*) d*z*, we can rewrite Eqn (B19) as:

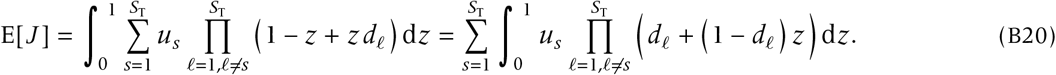

Eqn (B20) represents the general expression for the expectation of JD provided that species incidences are uncorrelated.

#### Break to check: experiments

Experiment 1 | When *S*_T_ = 1, we immediately get 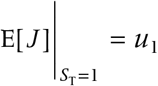. Thus the conditional expectation is *u*_1_ / (1 − *d*_1_).

Experiment 2 | When *S*_T_ = 2,

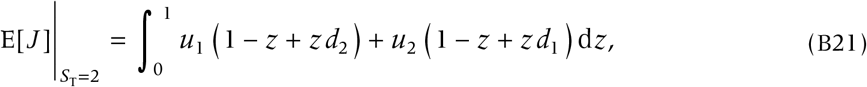

which is *u*_1_ (1 − 1/2 + *d*_2_ /2) + *u*_2_ (1 − 1/2 + *d*_1_ /2). Thus the conditional expectation is

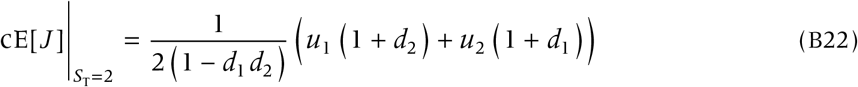

Experiment 3 | When all species are equal, that is when (*p*_1,*s*_, *p*_2,*s*_) = (*p*_*°*,1_, *p*_*°*,2_) with *p*_1,*s*_ *p*_2,*s*_ = *b* and *a*_1,*s*_ *a*_2,*s*_ = *d*,

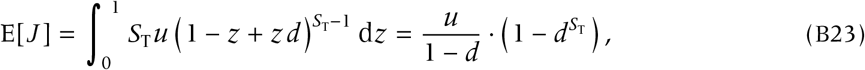

thus recovering Lu *et al*.’s (2019) results by dividing the RHS by 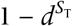.

#### Step 5: Use the beta function and elementary symmetric polynomials

For a real-valued vector 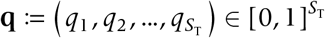, we denote the elementary symmetric polynomial of degree 𝓁 by σ 𝓁 (**q**), for 𝓁 = 0, 1, …, *S*_T_. For example, 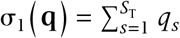, and 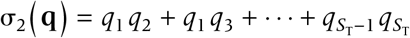. We also write **d**_*ŝ*_ for the vector (of length *S*_T_ − 1) that removes *d*_*s*_ from 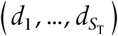, for *s* = 1, 2, …, *S*_T_. For example, 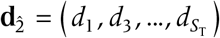.

Expanding the product in Eqn (B20) in terms of 1 − *z* and *z*, we get:

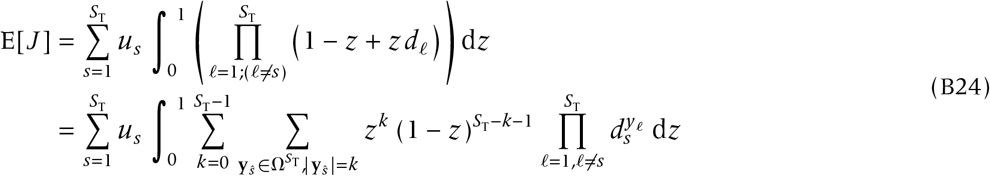

Using the Beta function 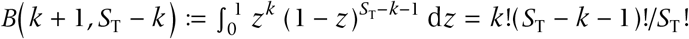, we can rewrite E[*J* ] as:

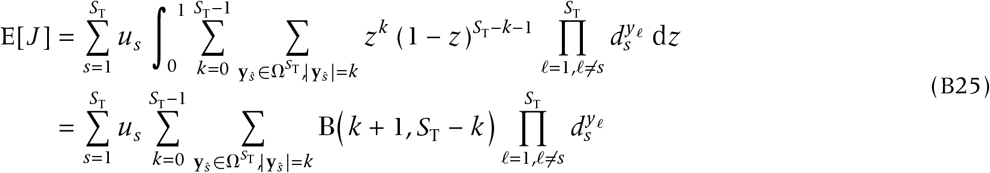

Using the *k*-th elementary polynomial σ_*k*_ (**d**_*ŝ*_) to replace the product term, we no longer need the summation over **y**_*ŝ*_ ; so:

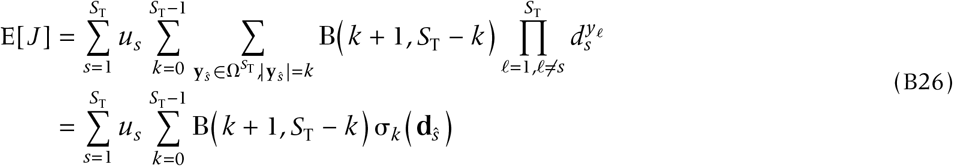

Finally, the conditional expectation is derived by dividing the unconditional expectation by 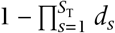. This form of the expectation may be more useful than the integral formula depending on computational environments.

#### More algebraic form

For mathematically inclined readers, we here provide more symmetric form of Eqn (B26). Notice the identity:

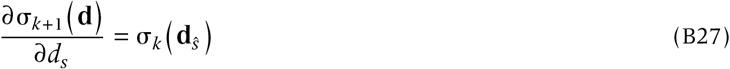

for any 1 ≤ *s* ≤ *S*_T_, 0 ≤ *k* ≤ *S*_T_ − 1. Substituting this into Eqn (B26) yields:

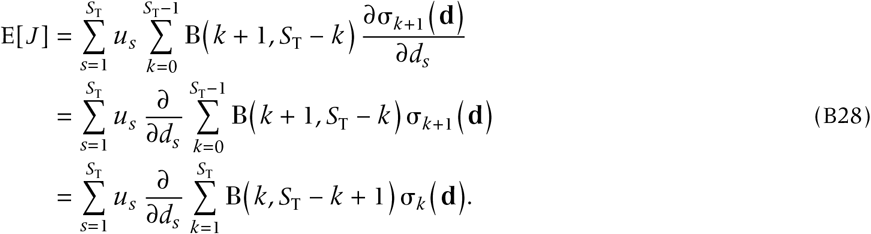

### Interpretation of the formula

Let us consider:

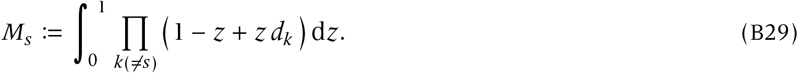

Using the identity of 1 − *z* + *z d*_*k*_ = *d*_*k*_ + (1 − *d*_*k*_) (1 − *z*), and changing the variable 1 − *z* ↦ *z*, we get:

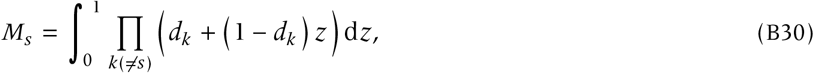

expanding which yields:

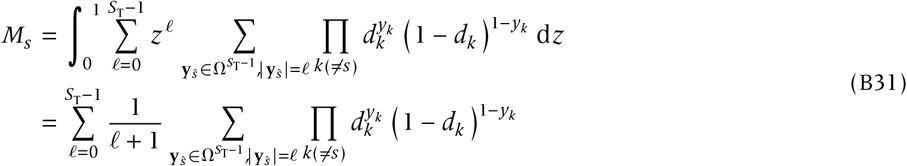

From this expression, the interpretation is clearer: suppose that the species *s* is unique; given this, we count the number *𝓁* of species present (excluding species *s*), considering all possible patterns of incidences of the other species 1, 2, …, *s* − 1, *s* + 1, …, *S*_T_, as is indicated by a binary sequence **y**_*ŝ*_ of length *S*_T_ − 1, where *y*_*k*_ = 1 if spcies *k* is double-absent, otherwise = 0. For instance, for *s* = 1 with **y**_î_ = (01101), species 3,4,and 6 are double-absent, whereas species 2 and 5 are not.

#### Examples

Given that species *s* = 1 is unique, which occurs with probability *u*_*s*_, we can obtain the probability that the other species *s* = 2, 3, …, *S*_T_ are either unique, common or double-absent. Inserting *S*_T_ = 2 yields *M*_1_ = (1 + *d*_2_) /2, because with probability *d*_2_, species 2 is absent from both sites, in which the contribution of species 1 to JD is 1, while with probability 1 − *d*_2_, species 2 is present, in which case the contribution of species 1 to JD is 1/2 (with species 2’s contribution not counted here), thus giving (1 + *d*_2_) /2. When *S*_T_ = 3, given that species *s* = 1 is unique, writing 00 for double-absence of species 2 and 3 and 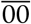 for non double-absence of species 2 and 3, we can write down all possible patterns each with a probability:

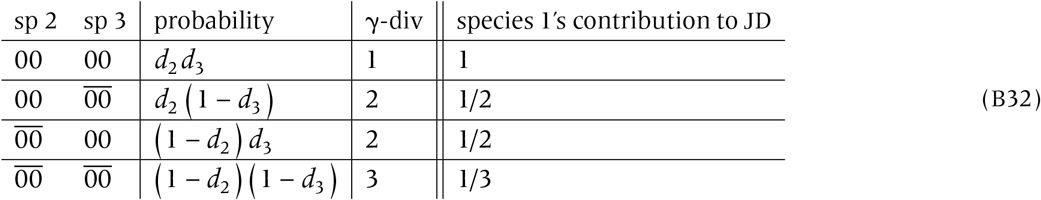

The expected contribution of species 1 to JD, conditioned on species 1 being unique, is thus given by

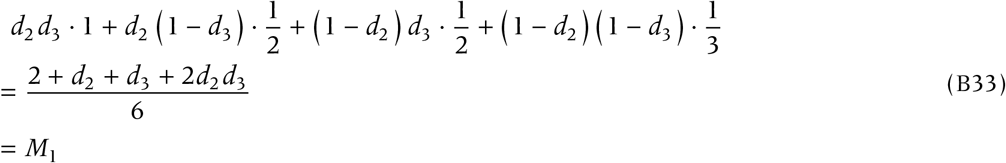

where the third line results from calculating the integral *M*_1_ for *S*_T_ = 3. From this reasoning, we can interpret Eqn (B20) as the sum of the conditional expectations of species’ contribution to JD.

### Heuristic approximation

#### Derivation

The conditional expectation can be obtained heuristically (Ontiveros *et al*. 2021):

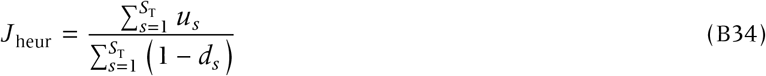

which represents the expected number of unique species divided by the expected number of present species. Deriving this formula requires quite a bit of calculations, but if we notice:

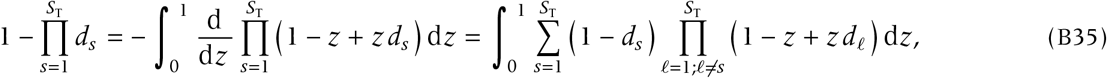

then we get:

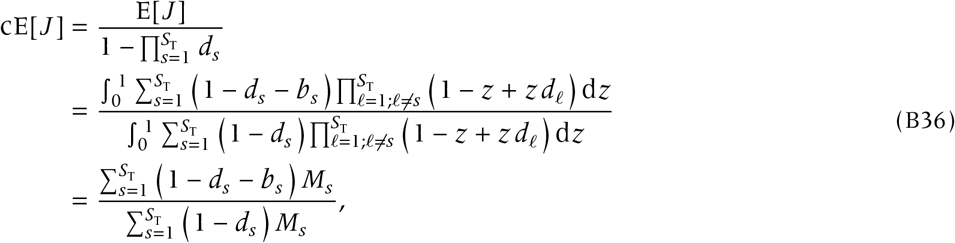

where we have put:

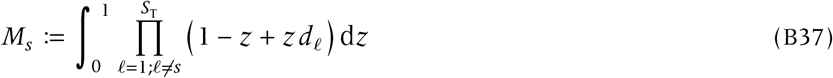

for *s* = 1, 2, …, *S*_T_. If we replace the integral *M*_*s*_ with 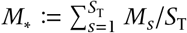 for *s* = 1, 2, …, *S*_T_, we then get:

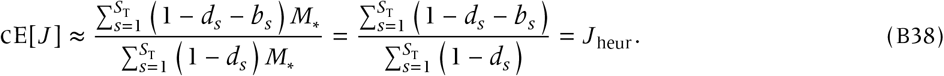

#### Asymptotic behavior

We may observe that *M*_*s*_ becomes increasingly smaller as *S*_T_ gets larger, where the asymptotic order is 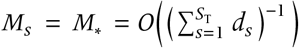, and thus, as *S*_T_ increases, the contribution of the replacement of *M*_*s*_ with *M*_*_ to the approximation error of the heuristic approximation becomes negligibly small, yielding the convergence of cE[*J* ] to *J* _heur_ (in probability).

#### Do flat SIPs yield a good approximation?

The heuristic approximation is, in a word, the proportion of the average uniqueness probability to the average probability of presence in at least one site. What if we instead consider:

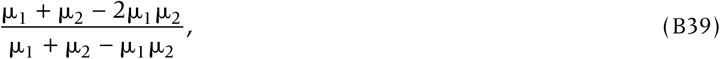

in which averages are take before taking the fraction? This question is equivalent to asking if the flat SIPs (assigning the average value to all species for each site) yield good approximations of the exact expectation of JD. We find that in this case, the error becomes systematically large (in absolute value; Figure S2). Jensen’s inequality may be applied to estimate inequality relationship between Eqn (B39) and *J* _heur_.

**Figure S2:**
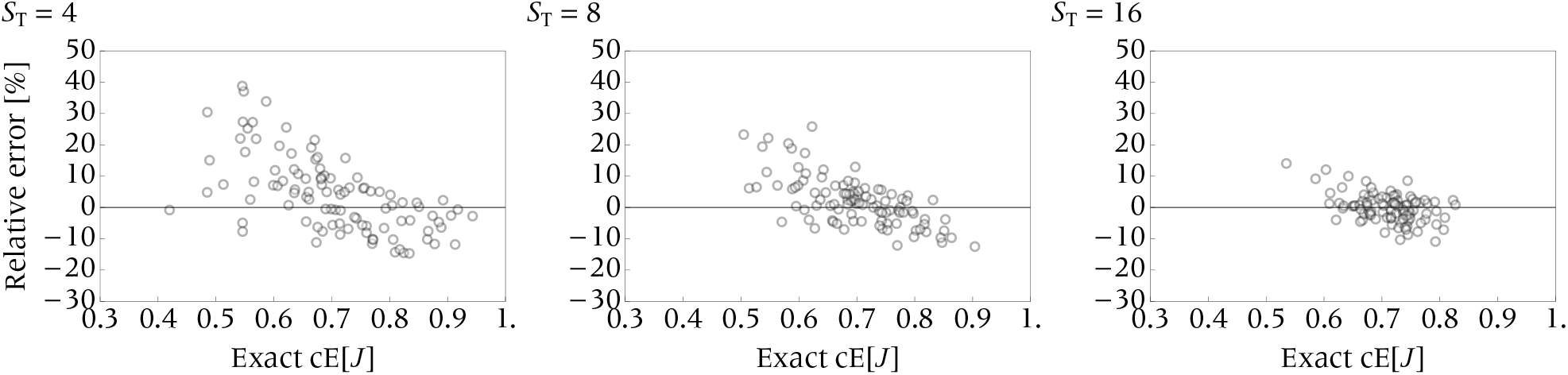
The relative error of Eqn (B39).

### Variance of JD

#### Same method as the mean

To compute the variance, we use the identity for a pair of positive quantities *v*_1_, *v*_2_ > 0:

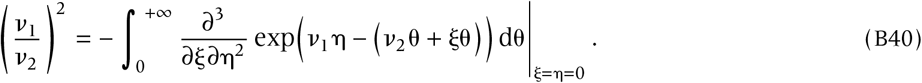

One may preferably differentiate the quantity before integration (otherwise, erroneous calculation is possible). For JD, we choose 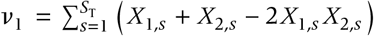, which represents the number of unique species, and 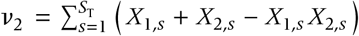, which represents the number of present species (gamma diversity). That is:

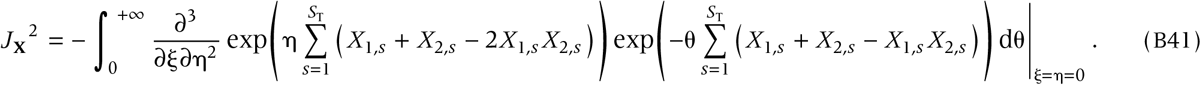

The expectation of *J*_**X**_^2^ is given by:

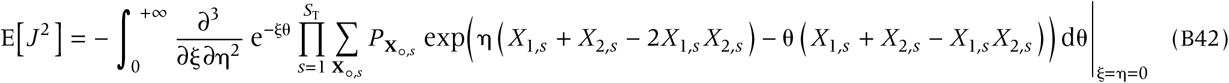

By evaluating the Boolean variable,

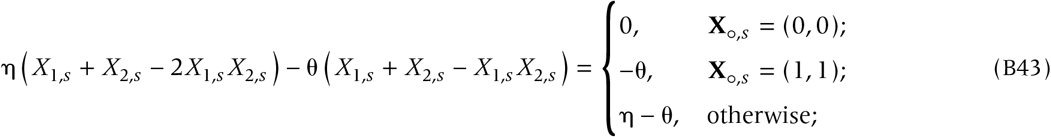

the resulting expression reads:

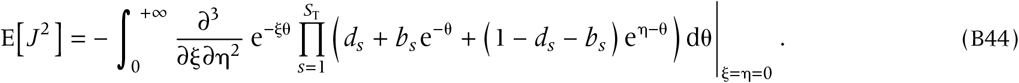

This is the most general expression for the second moment of the Jccard dissimilarity. For brevity we write τ 𝓁 (θ) := *d* 𝓁 + (1 − *d* 𝓁) e^−θ^ for the moment generating function of the probability that species 𝓁 is present in at least one of the sites, 1 − *d* 𝓁 ; write Ψ_*s*_ (θ, η) := *d*_*s*_ + *b*_*s*_ e^−θ^ + *u*_*s*_ e^η−θ^, so that Ψ_*s*_ (θ, 0) = τ_*s*_ (θ).

Leibniz rule for the second η-derivatives is given by:

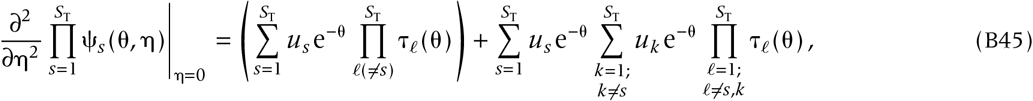

using which we get:

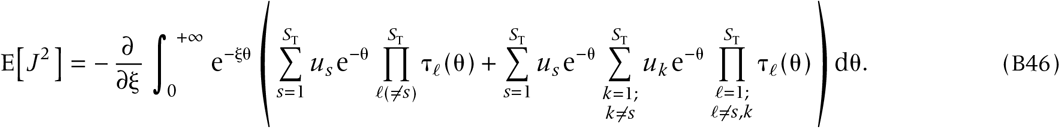

We can evaluate this integral as did we before. However, the resulting equation is heavily complicated (involving, e.g., the harmonic numbers) and computationally expensive.

#### Approximating variance using Hubbard-Stratonovich transformation

Here, we take a different approach to evaluate the variance. We use the identity:

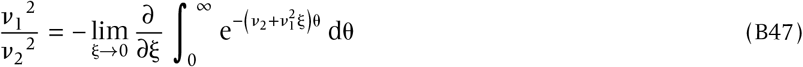

for 0 ≤ *v*_1_ ≤ *v*_2_, as well as the Hubbard-Stratonovich transformation (Hubbard 1959):

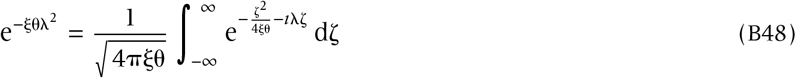

where *i* represents the imaginary unit. Combining the identities gives:

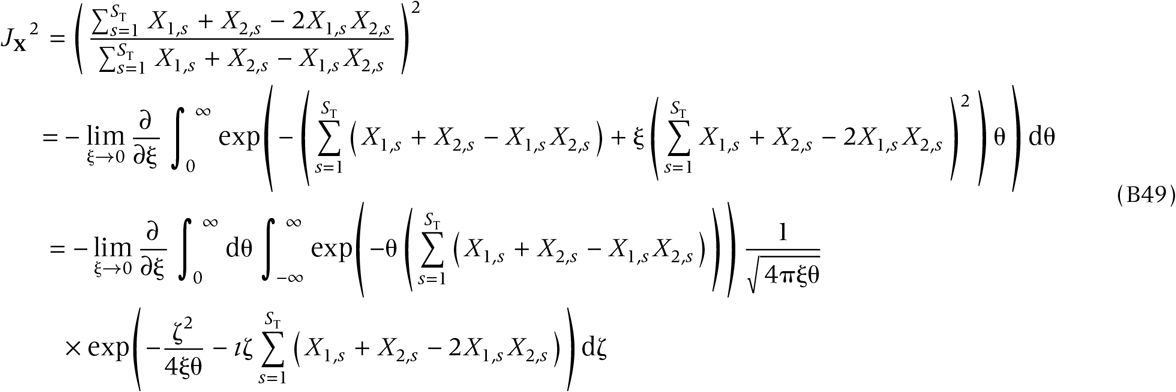

Let us evaluate the Boolean variable:

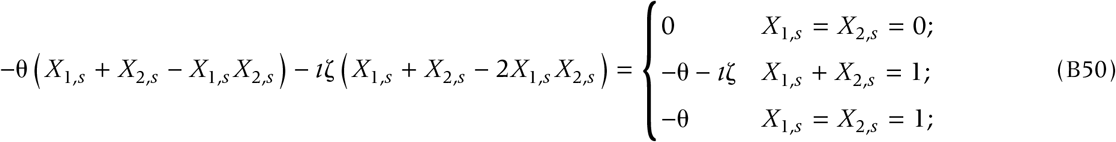

then we get:

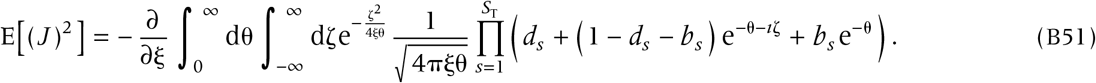

If we approximate the product as:

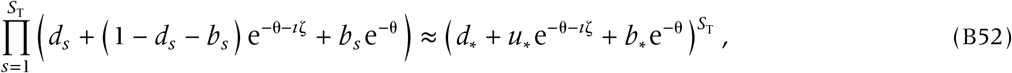

where the *-subscripted quantities are the arithmetic means, over *s* ∈ {1, …, *S*_T_}, of the corresponding quantities, i.e, 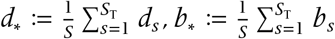, and *u*_*_ := 1 − *d*_*_ − *b*_*_, then the expected value is approximated by

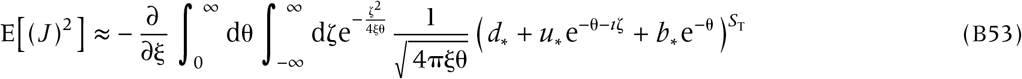

evaluated at ξ = 0.

Interchanging the order of the derivative and the double integral, we get

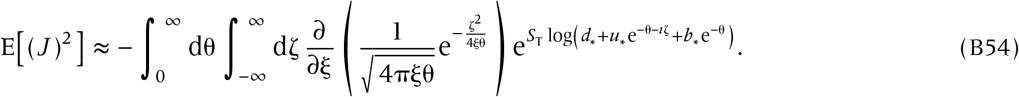

In the limit ξ → 0, the function 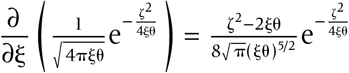 is very peaked about ζ = 0. Therefore, we expect the integrand to be nicely approximated if we substitute the logarithm by its series expansion about ζ = 0,

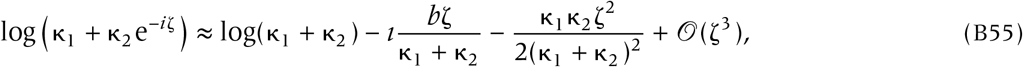

with κ_1_ := *d*_*_ + *b*_*_ e^−θ^ and κ_2_ := *u*_*_ e^−θ^. Inserting this second approximation into Eqn (B54) we get

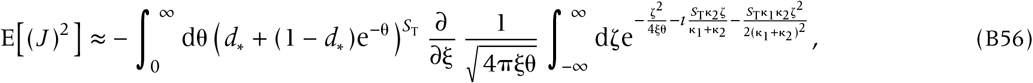

which, again, has to be evaluated at ξ = 0. The integral over ζ can be evaluated as

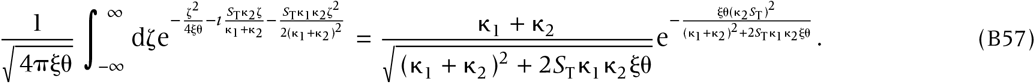

Now, we can take the derivative with respect to ξ and evaluate it at ξ = 0 to get

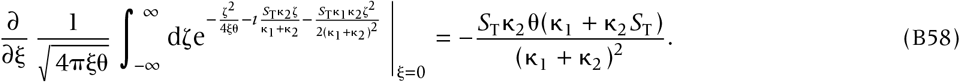

Therefore, inserting this expression into Eqn (B56) and replacing κ_1_ and κ_2_ by their expressions in terms of *d*_*_, *b*_*_, *u*_*_, and θ, we obtain

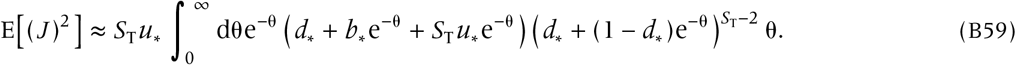

Changing to the variable *z* = *e*^−θ^ yields

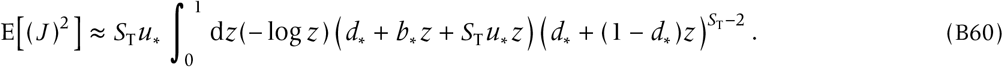

We now use the binomial expansion 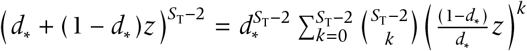 to get

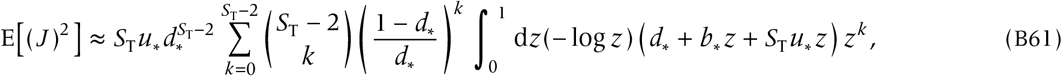

which, upon evaluation of the integral, yields

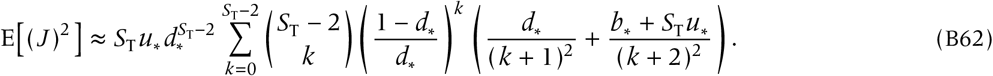

The sum above can be expressed in terms of generalized hypergeometric functions _*p*_ *F*_*q*_ ({*A*}, {*B*}; *Z*) as

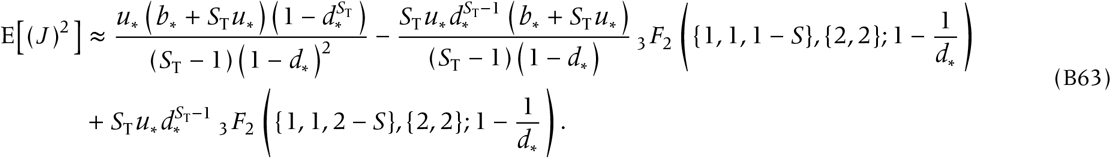

As a consequence, we find the following approximation for the variance,

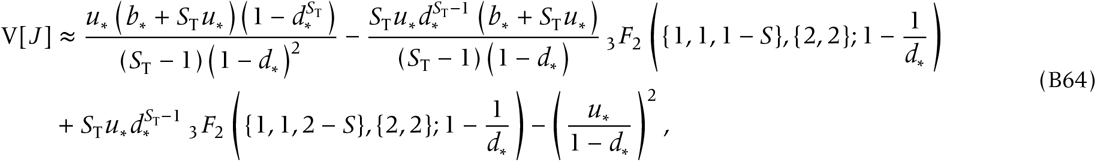

where we have approximated the expectation E[*J* ]^2^ with the square of our heuristic approximation,

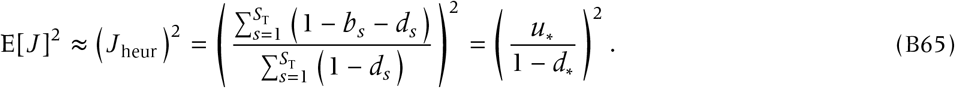

The analytical approximation obtained in Eqn (B64) yields always averaged standard deviation relative errors less than 10%. In most of the cases relative errors for the standard deviation, averaged over realizations of incidence vectors, are only about 2%.

#### Leading term in the limit of large *S*_T_

In order to get more insight about the dependence with *S*_T_ in the limit *S*_T_ → ∞, we have computed an asymptotic expansion of the variance to get the leading term in the series expansion on *S*_T_. First let us write Eqn (B59) as

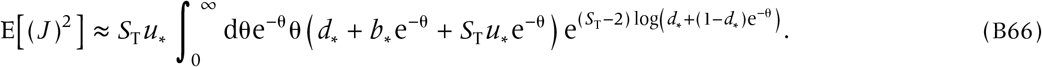

As *S*_T_ → ∞, the exponential function will be very peaked at the maximum of log (*d*_*_ + (1 − *d*_*_)e^−θ^). So we expect to have a good approximation in the limit *S*_T_ → ∞ if we replace the logarithm by its series expansion,

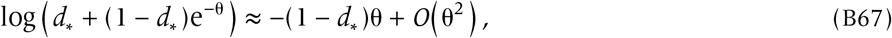

about the point at which the maximum is reached, i.e, θ = 0. Then, for large *S*_T_, Eqn (B59) will be nicely approximated by

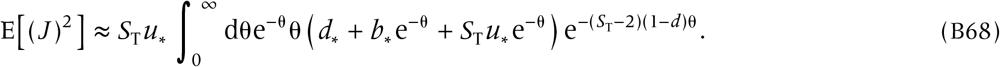

This integral can be actually evaluated to give

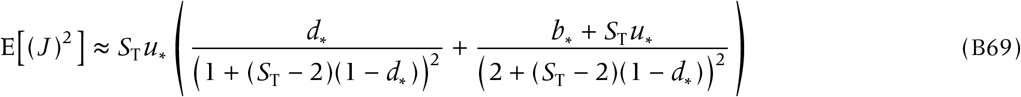

plus subleading terms in *S*_T_. Here we observe that our approximation for E[(*J*)^2^ ] converges to the squared heuristic Jaccard measure approximation,

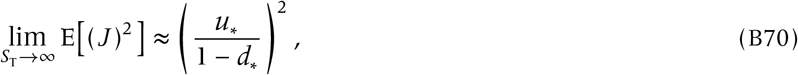

so, in the limit of large *S*_T_ we find the following leading term for the variance approximation:

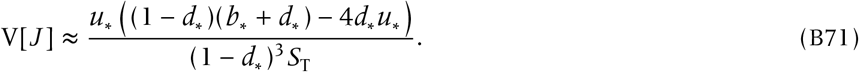

The variance decreases as *S*^−1^ in the case of large number of species. This explains why our heuristic approximation works very well in that limit.

## Appendix C Benchmark and key results

### Benchmark result

Eqn (B23) proves that when *p*_*r,s*_ = *p*_*r*, °_ for all *s* and for each *r* the heuristic approximation becomes exact for the conditional expectation of JD.

### Proof of key result 1, the transfer principle for beta

We here prove that, when *p*_1,*s*_ = *p*_2,*s*_ =: *p*_*s*_ for all species *s* = 1, 2, …, *S*_T_, the conditional expectation of JD decreases with increasing species unevenness in the presence probabilities, provided that 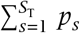 is fixed.

The proof results from the direct application of Schur’s condition as well as the symmetry property (Marshall *et al*. 1979, A.4.a and A.5, Chapter 3), so we set *S*_T_ = 2 and shall show the following:

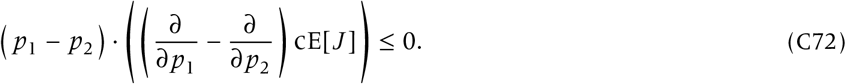

In the first place, the denominator 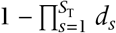 is Schur-convex, and as a consequence, its reciprocal is Schur-concave (Marshall *et al*. 1979, F.1.a). So we need to show:

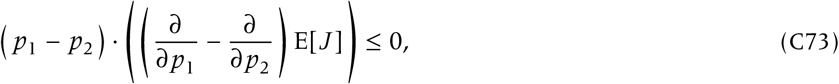

which follows from:

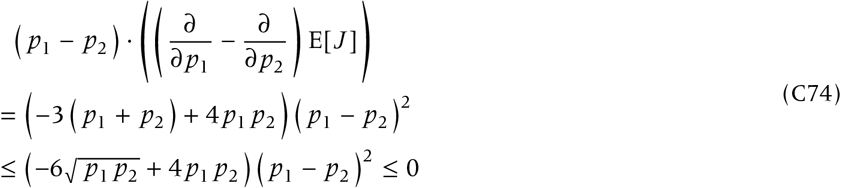

whenever *p*_1_, *p*_2_ ∈ [0, 1], completing the proof. Note, however, that this simple calculation is not applicable when the total presence probability can change; in that case, the notion of weak majorization may be needed.

### Numerical evaluation of the transfer principle for beta

To numerically check the transfer principle for beta, we define the following Hill function:

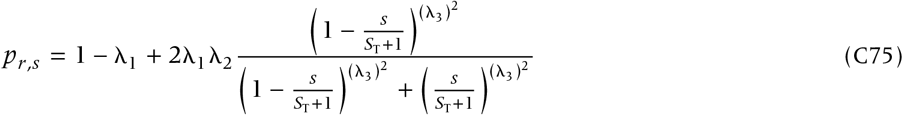

where the average presence probability is μ = λ_1_ + λ_2_ − λ_1_ λ_2_, and λs are parameters within the unit interval [0, 1]. The Hill coefficient, λ_3_, tunes the degree to which the curves are sloped at the intermediate species label. We choose *S*_T_ = 23, λ_1_ = 0.12, μ = 0.48 (with λ_2_ ≈ 0.41) and varied λ_3_ from 0 to 4 with a step size 4/50 = 0.08 along the horizontal axis. The expectation of JD is numerically evaluated using the expectation formula.

### Key result 2: two-species approach

#### Proof of Table 2

We here show the ‘Effect on JD’ column in Table 2.

Suppose that we have a given incidence matrix sized 2-by-*S*_T_, denoted **X**, and denote its JD by *J*. Specifically, we denote the number of (i) unique species for **X** by *U* or (ii) gamma-diversity by γ, implying *J* = *U* /γ.

We now append 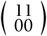 to **X**, to get a new incidence matrix denoted **X**′, and denote its JD by *J* ′. Similarly we write *U* ′ for the number of unique species for the incidence matrix **X**′ and γ′ for gamma-diversity by γ′: *J* ′ = *U* ′/γ′. Because *U* ′ = *U* + 2 and γ′ = γ + 2, we have:

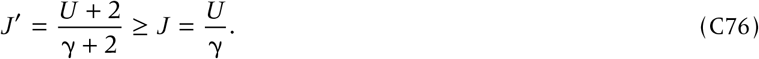

The equality occurs only if *J* = 1 (which is of limited interest). The increment in *J* is larger than appending 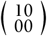 because of the addition of two unique species. Using the same argument gives the effects of 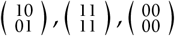 on JD. Finally, let us consider the effect of appending 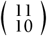 on JD. Using the analogous notation we get:

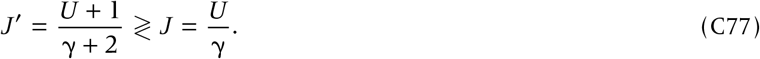

The inequality > (or <) occurs when *J* < 1/2 (or < 1/2, respectively), and the equality occurs when *J* = 1/2.

#### The commonality of histograms

We first show that all the fundamental SIPs have the same probability to generate the single-one incidence matrices. Let us write four values of presence probabilities *p*_1,*s*_ = *q*_1_ ≥ *q*_2_ ≥ *q*_3_ ≥ *q*_4_. The probability of the occurrence of single-one class is given by:

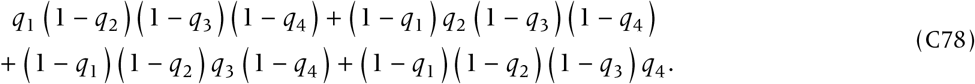

This quantity is symmetric for any permutations of (*q*_1_, *q*_2_, *q*_3_, *q*_4_), and thus is independent of reassignments of these values to *p*_*r,s*_. Indeed, the single-one class 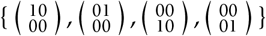 is closed with any permutations of the elements.

Analogously, single-zero, all-one, and all-zero matrices are fully symmetric, leading to the common presence probabilities.

#### Characterization of the fundamental SIPs: most likely incidence matrices

We here prove that, as a representative case, the anti-parallel nestedness SIPS and the parallel nestedness SIPs most likely produce the nestedness class of incidence matrices, but produce, with different probabilities, the checkerboard class.

The probability of the nestedness class of incidence matrices is given by:

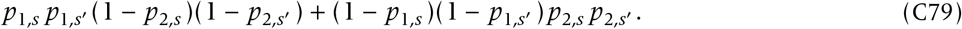

This probability is maximized when *p*_1,*s*_ and *p*_1,*s*_′ are the first and second largest, or when *p*_2,*s*_ and *p*_2,*s*_′ are the first and second largest (as can be easily checked), which occurs, respectively, for the anti-parallel nestedness or parallel nestedness SIPs.

The probability of the occurrence of the checkerboard class is given by:

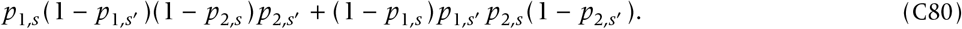

For the anti-parallel nestedness (or parallel nestedness) SIPs, it holds *p*_2,*s*_ < *p*_2,*s*_′ (or *p*_2,*s*_ > *p*_2,*s*_′, respectively), with *p*_1,*s*_ > *p*_1,*s*_′. Eqn (C80) is smaller for the latter SIPs (resulting from some arithmetic calculations), thus proving that the anti-parallel nestedness SIPs have a larger JD than do the parallel nestedness SIPs.

## Appendix D SDM

We use data of five woodpecker species, *Picus viridis, P. canus, Dendrocopos major, D. minor*, and *Dryocopus martius* in Switzerland (Schmid *et al*. 1998, 2018; Zurell *et al*. 2019b, 2020). These species have common evolutionary history but use different habitats (Benz *et al*. 2006; Pasinelli 2007; Pons *et al*. 2010). For example, *P. canus* and *D. minor* occur at lowlands, while *P. viridis* is more widely found across Switzerland. Since the variation in the estimated, geographic distributions reflects the incidence heterogeneity, the system is suitable for the analysis of temporal beta-diversity (Oliveira *et al*. 2023).

Data is collected over a four-year period (1993-1996) in usually three visits per year (2 above the treeline) using a simplified territory mapping approach, and integrates in the Swiss breeding bird atlas at 1-by-1 [km] resolution (Schmid *et al*. 1998, 2018). *The data source we have used includes environmental variables – climate, topography and vegetation structure – at the same spatial scale but geographic coordinates are removed. Data can be found at Zurell et al*. 2019b,a. We use the whole dataset to infer SDMs using only the climatic variables, as current and future values of these variables for Switzerland are available in worldclim (www.worldclim.org; Hijmans *et al*. 2005). We have downloaded current climatic data using function getData from the R package raster using argument name = ‘worldclim’, and future climate with the same function call with arguments name = ‘CMIP5’, rcp = 45, year = 50, and model = ‘NO.’ We have examined the degree to which woodpeckers’ metacommunity across Switzerland in 2050 will become dissimilar in the future climatic conditions (Representative Concentration Pathway 4.5; Van Vuuren *et al*. 2011) from the that under the current climate (spatial scale: 1 × 1[km]). For each species, we have used an ensemble approach of, initially, four different algorithms: generalized linear models (GLMs), generalized additive models (GAMs), random forests (RFs) and boosted regression trees (BRTs). However, GLMs and GAMs produce unreliable projections and are subsequently excluded from our analyses. We then project current and future incidences for each species in our ensemble approach. With those incidences, we calculate the expected dissimilarity (Eqn (3)) at each location of Switzerland for the subcommunity of woodpeckers.

**Figure S3:**
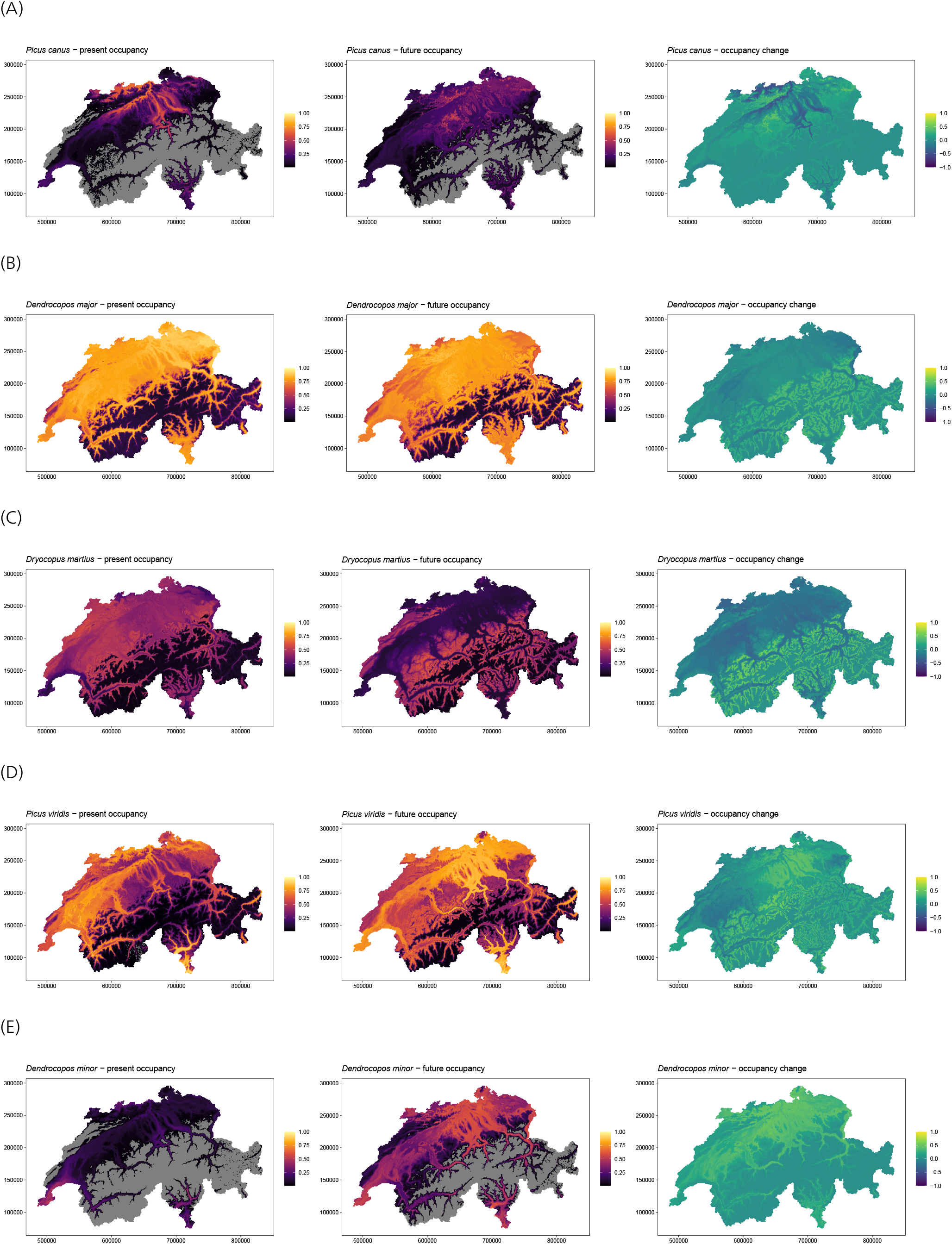
The presence probabilities: *p*_*s*,current_, *p*_*s*,future_, and *p*_*s*,future_ − *p*_*s*,current_.

